# Widespread occurrence of hybrid internal-terminal exons in human transcriptomes

**DOI:** 10.1101/2021.05.27.446076

**Authors:** Ana Fiszbein, Michael McGurk, Ezequiel Calvo-Roitberg, GyeungYun Kim, Christopher B. Burge, Athma A. Pai

## Abstract

Alternative RNA processing is a major mechanism for diversifying the human transcriptome. Messenger RNA isoform differences are predominantly driven by alternative first exons, cassette internal exons and alternative last exons. Despite the importance of classifying exons to understand isoform structure, there is a lack of tools to look at isoform-specific exon usage using RNA-sequencing data. We recently observed that alternative transcription start sites often arise near annotated internal exons, creating “hybrid” exons that can be used as both first or internal exons. To investigate the creation of hybrid exons, we built the HIT (Hybrid-Internal-Terminal) exon pipeline that systematically classifies exons depending on their isoform-specific usage. Using a combination of junction reads coverage and probabilistic modeling, the HIT index identified thousands of hybrid first-internal and internal-last exons that were previously misclassified. Hybrid exons are enriched in long genes with at least ten internal exons, have longer flanking introns and strong splice sites. The usage of hybrid exons varies considerably across human tissues, but they are predominantly used in brain, testis and colon cells. Notably, genes involved in RNA splicing have the highest fraction of intra-tissue hybrid exons. Further, we found more than 100,000 inter-tissue hybrid exons that changed from internal to terminal exons across tissues. By developing the first method that can classify exons according to their isoform contexts, our findings demonstrate the existence of hybrid exons, expand the repertoire of tissue-specific terminal exons and uncover unexpected complexities of the human transcriptome.

## Introduction

The composition of mRNAs in a eukaryotic cell is highly dynamic and diverse. Greater than 95% of multi-exonic mammalian genes express multiple mRNA isoforms and isoform usage is often regulated in a tissue and context-specific manner. Despite the recent focus on alternative splicing mechanisms, it was recently observed that isoform differences across human tissues are predominantly driven by alternative mRNA start and end sites (1–3). Alternative splicing of internal exons was estimated to explain tissue-dependent transcript differences for only ∼35% of the genes, while alternative transcription start and end sites accounted for the majority of tissue-dependent isoform usage (1). The modulated of transcripts through usage of alternative mRNA start and end sites is highly conserved during evolution, with alternative terminal exon usage occurring for most genes across yeast, plant, insect and mammalian species (4–6). Further, accumulating evidence suggests that a significant fraction of internal exons is used as first (FEs) or last exons (LEs) in different cell contexts (7, 8). The usage of polyadenylation sites in introns can lead to the conversion of an internal exon into a 3′ terminal exon (7) and cryptic promoters that arise during evolution may lead to conversion of internal exons to first exons (8). However, much less is known about the prevalence, usage, and evolution of this class of hybrid exons.

The usage of alternative transcription start and end sites can contribute to the regulation of the transcriptome and proteome in many ways. Unlike other exon types, relatively few alternative terminal exons directly code for alternative protein sequences. Instead, alternative first or last exons are more likely to influence the open reading frame through introduction of alternative start codons or induce protein truncation with premature stop codons. However, the vast majority of alternative terminal exons contribute to alternative 5’ or 3’ untranslated region (UTR) composition and consequent regulation of post-transcriptional mRNA processes (9–11). Although UTRs do not directly contribute to protein sequence, they harbor crucial regulatory sequences (including those for RNA-binding proteins and miRNAs (12)) and are thus modulators of mRNA subcellular localization (13), stability (14) and translation (15) in vivo. Unsurprisingly, given the many ways that alternative terminal exons can influence mRNA and protein diversity, the dysregulation of transcription start and end sites has been associated with several human diseases, including cancers, and neurological disorders (16, 17). Further, the usage of alternative terminal ends are dynamically regulated after exposure to environmental or immune stimuli (18, 19), suggesting that these mechanisms may also play an important role in cellular responses and remodeling. Given the central role of terminal exon composition in the regulation of both the transcriptome and the proteome, it is crucial to be able to properly classify which exons are used as terminal or internal exons across cell types and cellular contexts.

Many experimental strategies have been developed to specifically identify transcription starts and ends by anchoring on the 5’ 6-methyl-guanosine cap (ie. CAGE-seq) or the 3’ polyA-tail (ie. 3p-seq). However, these experiments have limited utility, since they are unable to simultaneously quantify expression of mRNA molecules or provide information about the isoforms associated with the terminal site usage. Furthermore, these specialized protocols are laborious, making them less practical than RNA-sequencing (RNA-seq) for general use across cell types and cellular conditions. Given the widespread use and availability of RNA-seq datasets, there is significant interest in using RNA-seq data to identify and quantify the usage of alternative terminal events (20–23). However, despite many efforts in the past few years, identifying terminal exons from short-read transcriptomic alignments and quantifying their usage remain challenging. Of specific interest is how to uniquely classify exons as first, internal and last, which we refer to as the unsolved “exon type problem”. These issues are especially problematic for hybrid exons, which would be classified as internal in RNA-seq data and terminal in data from the specialized terminal end sequencing techniques highlighted above, despite being used as both terminal and internal exons in different transcripts within a cell type. Even the newest classification and quantification algorithms (23–25) are not able to differentiate between purely terminal exons and exons with hybrid features, let alone quantify the proportional usage as a terminal or internal exon.

Here, we introduce the Hybrid-Internal-Terminal (HIT) index, developed to systematically identify and classify exons as hybrid, internal, or terminal based on short-read RNA-seq data. By modeling the ratio of exon-exon splice junction reads (SJRs), the HIT index pipeline reliably classifies exons and identifies thousands of hybrid first-internal or hybrid internal-last exons. By comparing HIT index exon classifications with classifications from specialized techniques to identify transcript ends, we show that our approach accurately classifies exons based on their isoform-specific usage. After applying the HIT index to RNA-seq data across human tissues, we are able to characterize features that differentiate hybrid terminal exons. Overall, the widespread identification of hybrid exons expanded the repertoire of tissue-specific isoforms and highlighted the diversity of the human transcriptome.

## Results

The regulation of mature RNAs can occur at multiple levels within cells, including quantitatively by varying gene expression levels and qualitatively by varying isoforms through alternative transcription start site, splice site, and/or cleavage site usage. Advances in high-throughput RNA-seq have provided unprecedented insights into mRNA levels across cell types, the internal structure of isoforms, and novel alternatively spliced exons. However, it is still challenging to use short-read RNA-seq data to also identify novel terminal ends of mRNAs - specifically, 5’ transcription start sites and 3’ transcription end or polyadenylation sites - or quantify their alternative usage. This is particularly difficult for hybrid terminal exons, which can be used as either terminal and internal exons in the same cell type. The ability to measure terminal site usage simultaneously with mRNA levels and isoform usage would enable a complete picture of the dynamics of both alternative untranslated regions and internal coding exons. To overcome these challenges, we set out to develop an approach that robustly identifies and quantifies hybrid exons, internal exons, and exons used only as terminal exons.

### The HIT index classifies exons based on their isoform-specific usage

We developed the Hybrid-Internal-Terminal (HIT) index, the first metric that leverages informative SJRs in short read polyA RNA-seq data to classify exon positioning and usage within an isoform. SJRs in RNA-seq data can be used to discern the connectivity between two exons that have been spliced together. Internal exons should have approximately the same number of SJRs connected to both upstream and downstream flanking exons. On the other extreme, the SJR distribution around terminal exons should be skewed such that there are only junctions connected to gene-proximal flanking exons (downstream flanking exons for FEs, or upstream flanking exons for LEs,). Thus, we rationalized that hybrid exons should have an intermediate distribution between FE and internal, or between internal and LE, with both upstream and downstream SJRs present, but a skew towards more gene-proximal SJRs depending on the terminal positioning of the hybrid exon. Building on these assumptions, the HIT index calculates the bias in upstream and downstream SJRs for each annotated exon (Figure 1A). The HIT index pipeline involves 3 primary steps: (1) annotating meta-exons from a transcript annotation, (2) extracting overlapping upstream and downstream SJRs for each exon, and (3) calculating the HIT index, classifying exons, and estimating percent spliced in values for alternative terminal exons. Since exact exon boundaries are confounded by imprecision in or alternative transcription start sites (TSS), 3’ and 5’ splice sites, and/or transcription end sites (TES), we use non-redundant metaexons (generated by collapsing all overlapping annotated exons) with an added buffer region. Statistical significance for the HIT index metric is estimated using a bootstrapped confidence interval and a measure of the probability that the HITindex was within ±10% of the observed value (Methods). With this formalization, exons used solely as FEs or LEs have a HIT index of –1 or 1, since they only have downstream or upstream SJRs, respectively. Internal exons, which should have roughly equal numbers of upstream and downstream SJRs, tend to have a HIT index near 0. Notably, this approach allows us to classify hybrid exons, which exhibit |HIT| indices between 0 and 1 commensurate with the fraction of transcripts that include a hybrid exon as a terminal or internal exon.

**Fig. 1.**
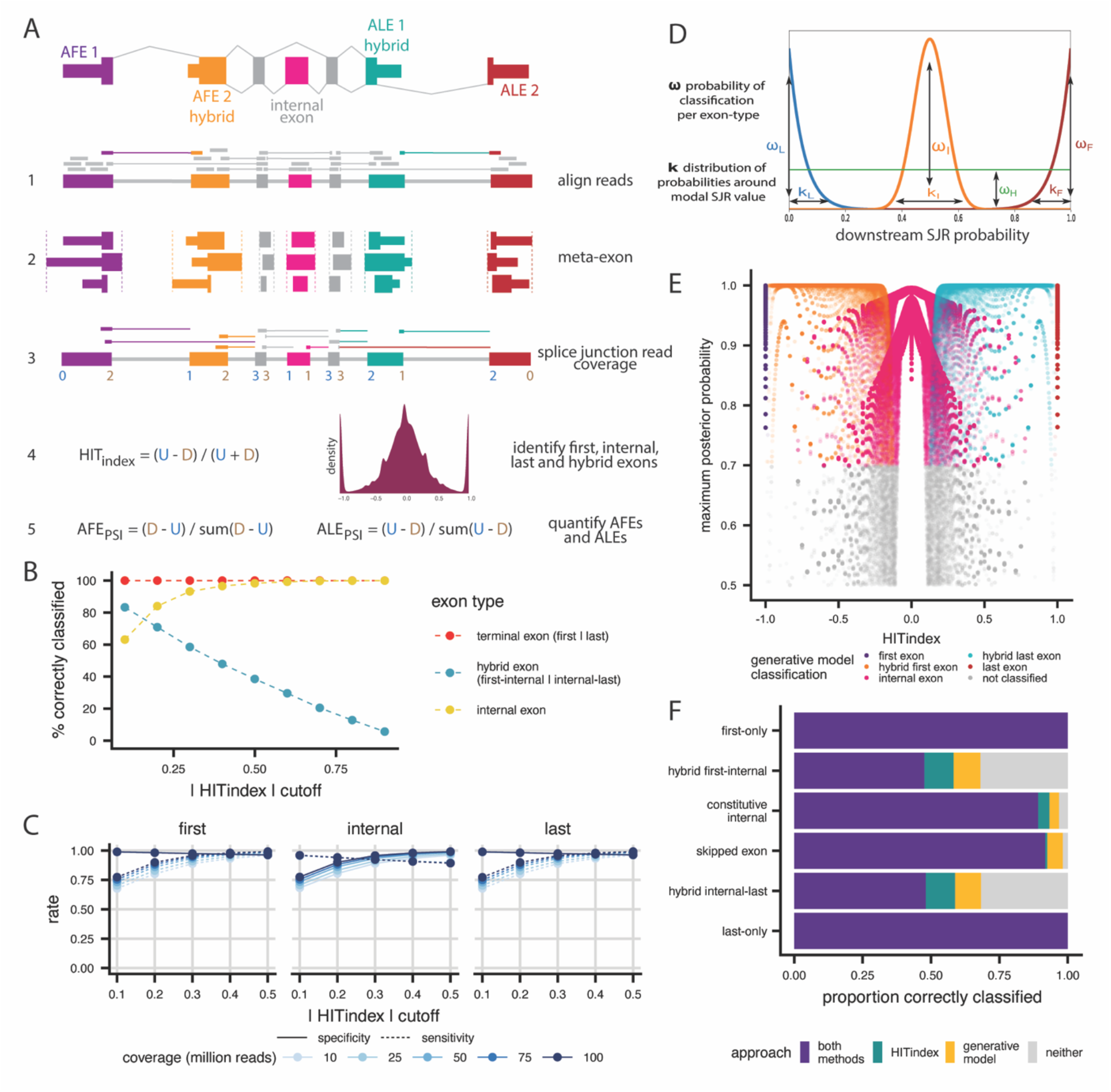
HITindex performs well with simulated data. (**A**), Schematic of the steps involved in the HIT index pipeline, including (1) alignment of reads to (2) meta-exons created by collapsing overlapping annotated exon boundaries, (3) extracting SJRs, and calculating (4) HIT index and (5) PSI values for each exon. (**B**), Percent of simulated exons correctly classified (y-axis) within each exon-type across variable HIT index thresholds (x-axis). (**C**), True positive (sensitivity, dotted lines) and true negative (specificity, solid lines) rates (y-axis) for simulated exon classification across moderate HIT index thresholds (x-axis), for different read depths. (**D**), Schematic of the mixture of beta-binominal distributions of downstream SJR counts (x-axis) that represent the generative model probabilities (y-axis) that an exon belongs to a particular exon-type, where the modes of the distributions are 0, 0.5, and 1 for last-only, internal-only, and first-only exons, respectively and hybrid exons are those that fall outside these distributions. (**E**), The relationship between the HIT index metrics (x-axis) and the maximum posterior probability from the generative model (y-axis) for various simulated exon types. (**F**), The proportion of different exon types that are correctly classified by only the HIT index, the generative model, both models, or neither.

### HITindex accurately classifies simulated exon data

To assess the performance of our approach, we simulated mRNAs with a range of gene structures (varying numbers, usage, and lengths of terminal, hybrid, alternatively spliced, and constitutive exons) and gene expression levels, and created short reads using a range of library preparation parameters, including different fragment sizes and read lengths (Figure S1, Methods). We then calculated the HIT index for each simulated exon using exon-exon junction reads as described above and classified exons as terminal exons that serve solely as FEs or LEs in a transcript, internal exons, and hybrid exons that can be used as either terminal or internal exons in a transcript. Classifications were performed as described in the Methods, with variable HIT index thresholds when specified. Overall, HIT index classifications recapitulate expected exon classifications at loose to moderate HIT index thresholds that represent less than a 50% imbalance (|HIT index| < 0.5, Figure 1B). Conversely, the classifications capture a smaller portion of hybrid exons at stringent thresholds of greater than 50% imbalance (|HIT index| > 0.5), but with higher confidence that each exon is truly hybrid. Since terminal-only exons entirely lack gene-distal junction reads and thus have extreme HIT indices of 1 or –1 (for FEs or LEs, respectively), all terminal exons can be reliably classified regardless of the chosen threshold. However, there is a tradeoff between the identification of hybrid exons and internal exons, where more hybrid exons are correctly identified at less stringent thresholds at the expense of the proper identification of internal exons.

We next sought to use these simulation data to identify systematic biases and characteristics of the data in situations where the HIT-index is unable to correctly classify hybrid or internal exons. The sensitivity for identifying terminal exons increased with more stringent |HITindex| thresholds, coupled to slightly reduced specificity in terminal exon classification (Figure 1C). Given that >95% of terminal-only exons (Figure 1B) are correctly classified, the variance in predictive values is entirely driven by hybrid exon classification. Both the false positive and false negative rates of terminal exon classification are less than 10% when using |HITindex| ≥ 0.3. However, the increased ability to correctly classify terminal exons across |HITindex| thresholds occur at the expense of the sensitivity, but not specificity, of internal exon classification. In both cases, overall specificity and sensitivity scale with the read depth of libraries (Figure 1C) and the minimum number of SJRs required (Figure S2A). Based on these trends, we further probed the parameters that influenced internal exon and hybrid exon classification across the chosen thresholds.

Exons that are solely used as internal exons should have an even distribution of upstream and downstream junction reads, however, we see that as many as 40% of internal exons are mis-classified at less restrictive |HITindex| thresholds, with as much as a 30% imbalance of junction reads. We reasoned that mis-classification of internal exons might be associated with their position in the gene, and therefore evaluated the mis-classification of exons stratified by their order in the gene (first, second, or third internal exon), the length of the exon, and the lengths of the upstream or downstream flanking exons (Figure S2B,C). Internal exons are more likely to be mis-classified as either FEs or LEs when the length of upstream or downstream exons, respectively, are shorter. Further, this mis-classification is more extreme when the library has larger insert sizes (Figure S2B) or is sequenced with shorter read lengths (Figure S2C). These trends suggest that our ability to classify internal exons that are near terminal exons is affected by RNA-seq edge effects, whereby there is a depletion of reads at the edges of molecules caused by size selection to achieve a narrow distribution of fragment lengths before sequencing. We do not see these same trends for skipped exons (SE), which were always flanked by two constitutive exons in the simulations. Instead, the ability to classify an SE as an internal exon is dependent on the percent spliced in (PSI) of the skipped exon, where lower SE inclusion results in a moderately higher frequency of mis-classification (< 10%) due to decreased junction read coverage (Figure S2D). To account for constitutive internal exon mis-classification, we incorporated an edge effect flag, which uses piecewise regression to identify exons that are likely to be affected by the edge effect (Figure S1C, Methods) and allows the end user to decide whether to include or exclude those exons from downstream analyses.

Since the HIT index represents the imbalance between upstream and downstream junction reads, the choice of HIT index threshold has a large impact on the ability to identify hybrid exons. To assess this impact, we stratified the percent of hybrid exons correctly classified by HIT index thresholds and the percent usage of the hybrid exon as a FE, measured by the percent spliced in (PSI) value (Figure S3A). Regardless of HIT index threshold, moderate to minimal usage of the hybrid exon as a FE rather than an internal exon reduces the ability to classify the exon as a hybrid exon, because junction reads are less skewed. At more stringent thresholds (|HIT index| > 0.5), the detection of hybrid exons with an FE PSI < 0.5 drops below 25%. When we evaluate the classification of hybrid exons at a moderate HIT index threshold of 0.3, this relationship becomes even more evident, where the assignment to hybrid decreases as PSI decreases and these exons are classified as solely internal exons.

To alleviate these issues, we developed a probabilistic framework to classify exons by directly modeling the probability of any exon being a hybrid, internal, or terminal exon considering read coverage and the ratio of downstream junction reads to total junction reads (Methods, Figure 1D). Specifically, this generative approach relies on jointly modeling the ratio of downstream junction reads to all junction reads for first, internal, and last exons, assuming that these exons are derived from probability distributions with a mode of 1, 0.5, and 0, respectively, with variance among the different exons for each class. Thus, the downstream junction read probabilities can be modeled as arising from a mixture of these three distributions and exons can be assigned a posterior probability of belonging to each distribution. Hybrid exons are detected as outliers relative to each of the non-hybrid distributions, enabling the estimation of a posterior probability of being either a first-internal or internal-last hybrid exon. Importantly, this method accounts for the intrinsic noisiness in sequencing data and the overdispersion of read counts relative to simple binomial or Poisson count distributions. By moving away from a strict HIT index threshold and allowing for flexibility in the classification probability of an exon, this framework enables exons to be correctly classified across the distribution of HIT indexes (Figure 1E). Most exons, including all terminal exons, are correctly classified by both the HIT index and generative model (Figure 1F). However, combining the two approaches leads to an increase in the classification of hybrid and internal exons, where the HIT index has more power for exons with lower read coverage, while the generative model has more power to classify hybrid exons with moderate PSI values (Figure S3B). All together, the combined approach is able to correctly classify 68% of simulated hybrid exons and 97% of internal exons.

### Identification of hybrid exons in human transcriptomes

We applied the HIT index pipeline to a high-coverage human RNA-seq data from LCL cells to evaluate the performance of our new method in classifying bonafide terminal, internal and hybrid exons. From a total of 353,677 annotated metaexons, we discarded 50% that are not expressed and 0.3% located in single-exon genes (Figure 2A). Using an absolute HIT index cutoff of 0.3 classified ∼60,000 internal, ∼16,000 terminal and ∼30,000 hybrid exons. (Figure 2B). We divided hybrid exons into hybrid FE-internal and hybrid internal-LE, each corresponding to ∼13% of total expressed exons (Figure 2C). Consistent with the simulated data, metaexons classified as hybrid by the HIT index have significantly higher probability of being classified as hybrid by the generative model (Figure 2D,E). Thus, we divided hybrid exons into high, medium or low confidence classifications depending on the bootstrapping variance metric (Methods, Figure 2F) and estimated the probability of consistent classification by the generative approach. As expected, the HIT index has lower power to identify hybrid exons relative to exons solely used as internal or terminal (Figure S4A,B). to study the properties of hybrid exons, below we consider only high confidence hybrid exons, which we define as being classified as a hybrid using the HIT index and a first-internal or internal-last posterior probability greater than 0.5 when using the generative model (Figure 2G). With these conservative criteria, we can reliably classify 3.82% of exons as hybrid in this dataset (Figure S4C).

**Fig. 2.**
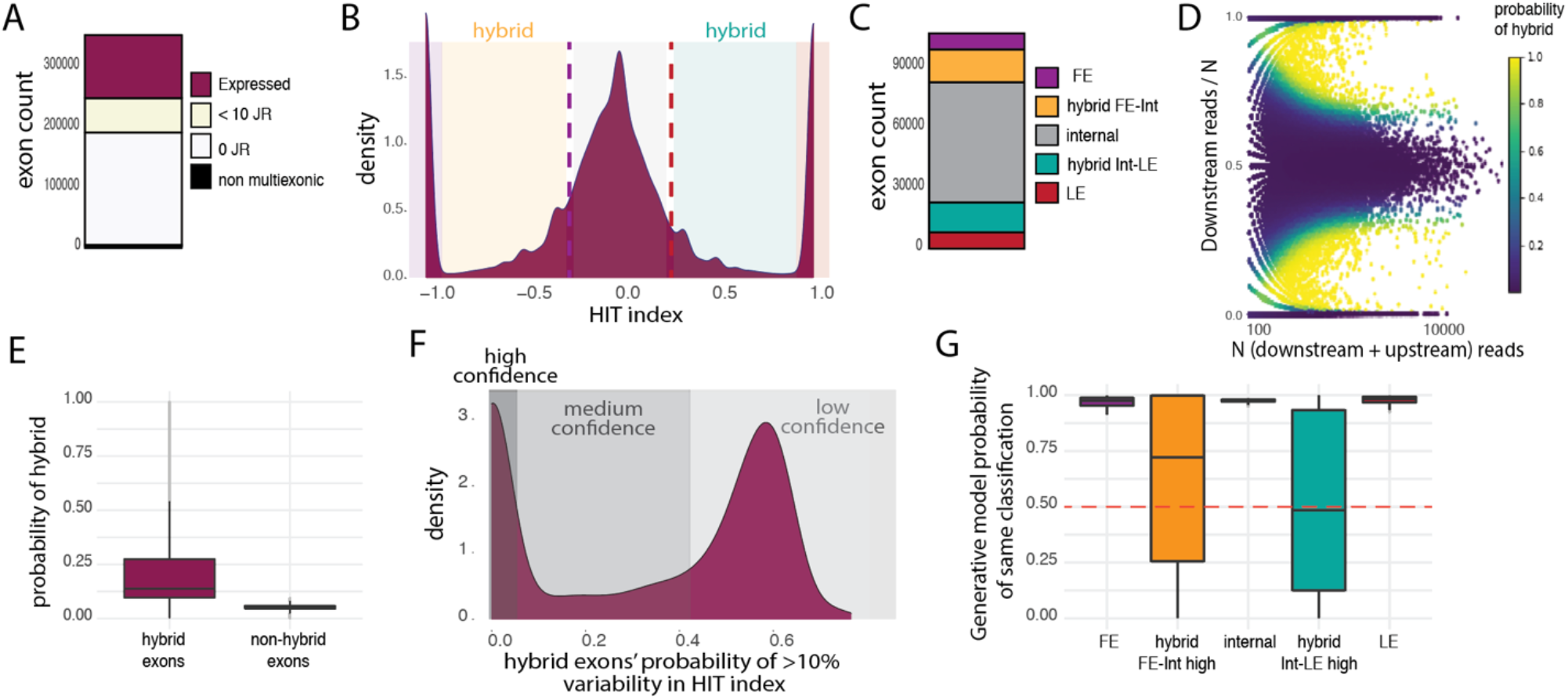
Classifying hybrid exons in RNA-seq data. (**A)**, The number of exons in multiexonic genes expressed in human lymphoblastoid cell lines (LCLs). (**B)**, Distribution of HIT indices (*x-axis*) for expressed LCL exons (> 10 junction reads). (**C)**, The classification of expressed exons into each of the 5 exon-types including both hybrid categories. (**D)**, Distribution of the ratio of downstream SJRs to total (N) SJRs (*y-axis*) relative to N (y-axis) across expressed exons. Heatmap represents the probability that an exon is classified as a hybrid exon by the generative model. (**E)**, Probability that an exon is classified as hybrid by generative model (*y-axis*) for exons classified as hybrid or non-hybrid by the HIT index metric alone (*x-axis*). (**F)**, Distribution of probability of >10% variability in HIT index (*x-axis*) for hybrid exons, binned in three confidence intervals. (**G)**, Probability of being classified in the same category by the HIT index metric and the generative model (*y-axis*) for all 5 exon-types (*x-axis*).

### Reference techniques validate HIT index classifications

In order to evaluate the accuracy of the HIT index classifications from RNA-seq data with orthogonal datasets, we used CAGE (26) and 3’end sequencing (27) datasets for direct assessments of TSS and TES usage, respectively. All HIT index classified FE and hybrid FE-internal exons overlapped with CAGE-identified TSSs, while all LE and hybrid Internal-LE exons overlapped with 3’sequencing-identified TESs (Figure 3A, B). Reassuringly, terminal and hybrid exons classified as first or last did not significantly overlap the TESs or TSSs, respectively, supporting high specificity of HIT index classifications. Moreover, neither gene expression thresholds nor |HIT index| cutoffs change TSS and TES specificity significantly (Figure S5). Overall, these orthogonal reference techniques validated HIT index classifications and demonstrated 100% specificity in determining true positive TSS and TESs. The HIT index also reliably identified hybrid exons, significantly expanding the landscape of exons that can act as FE and LE.

**Fig. 3.**
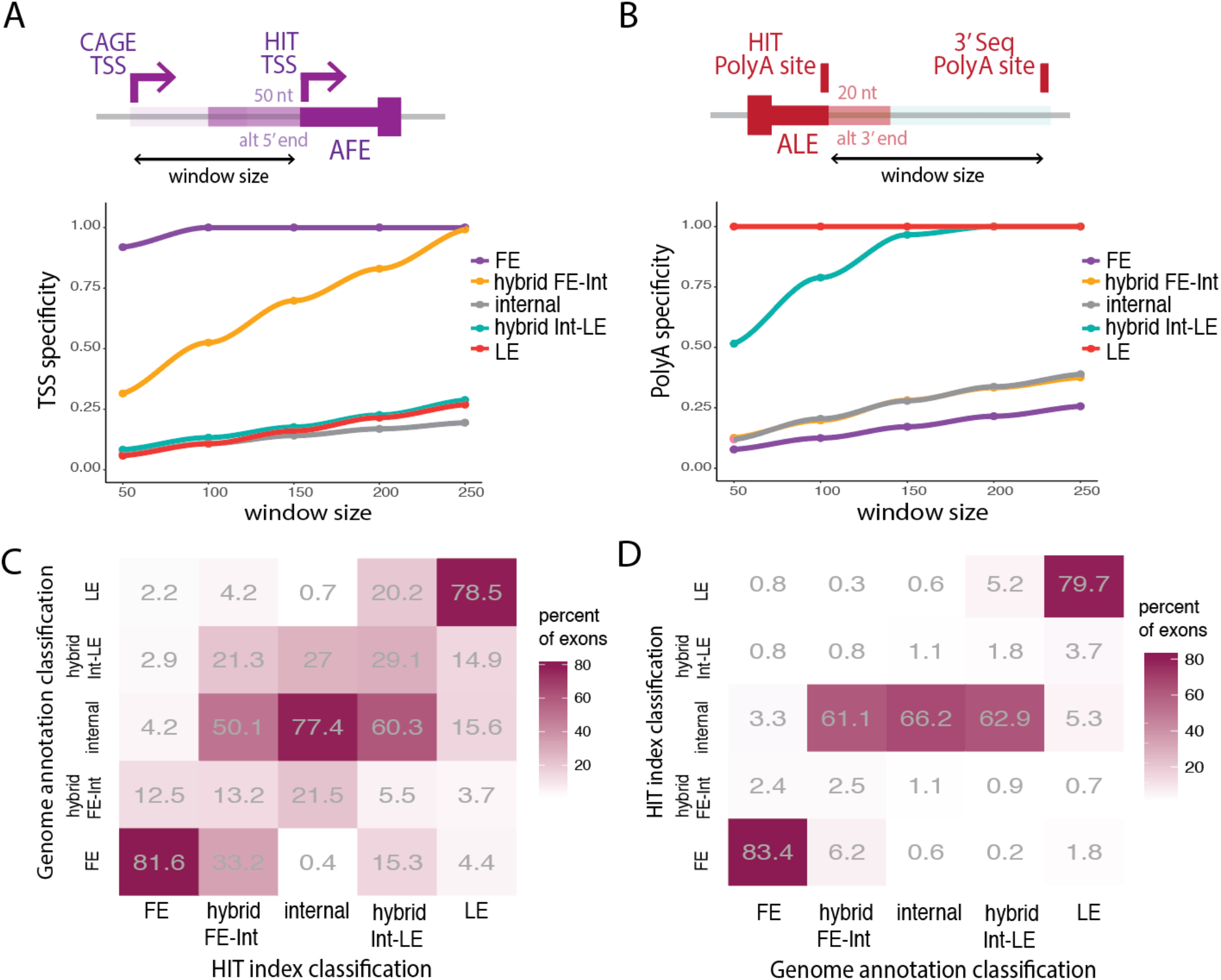
Evaluation of HIT index performance. TSS specificity. (**A**) and polyA specificity (**B**) for all five exon-types across window sizes, where specificity is calculated as the fraction of HIT index TSSs (start coordinate of identified FE) or HIT TESs (end coordinate of identified LE) that fall within the given distance away from any CAGE-seq identified TSS or 3’seq-identified polyA site. (**C)**, Heatmap of the percent of exons classified by the HIT index that are annotated as belonging to the same or different categories in the latest genome annotation. (**D**), Heatmap of the percent of exons annotated in the latest genome annotation that are classified into the same or different category by the HIT index.

We then compared HIT index classifications to annotated exon classifications (see Methods). While most HIT index terminal and internal exons were annotated as the same, the majority of hybrid exons were solely annotated as internal exons in genome annotations (Figure 3C). This suggests that current genome annotations underestimate the existence of hybrid exons. A similar underestimation of hybrid exons is true when conditioning on annotated exons and comparing to HIT index classifications (Figure 3D). Consistent with previous work (28), we find that only a minority of annotated exons are expressed in a single cell type and of these, many hybrid exons are mis-classified as internal exons. Together, our new classifications show that a reliance on genome annotations in RNA-seq analyses underestimates the usage of alternative terminal exons by mis-classifying hybrid exons.

We also used Oxford Nanopore direct RNA long-read sequencing data from LCLs to validate the HIT index classifications in selected genes that we highlight here (Figure 4). First, SFPQ is a member of the proline/glutamine-rich family of splicing factors and is associated with amyotrophic lateral sclerosis (29) Alzheimer’s disease and frontotemporal dementia (30). The tenth exon of *SFPQ* is used as a terminal exon in the longest isoform of *SFPQ* and is essential, since isoforms that skip exon 10 are non-coding or degraded by NMD. When exon 10 is excluded, splicing switches to a group of exons in the 3’-UTR, with exon 14 used as an alternative LE. We found that exon 12 is a hybrid FE-internal exon, used as an internal exon in transcripts that skip only exon 10 and as an FE in transcripts that skip exon 10 and use exon 14 as a LE (Figure 4A). SFPQ binds its own mRNA (Figure 4B), suggesting self-regulation of alternative splicing and polyadenylation (31). This suggests that the hybrid usage of *SFPQ* exons could contribute to the modulation of levels of active SFPQ protein.

**Fig. 4.**
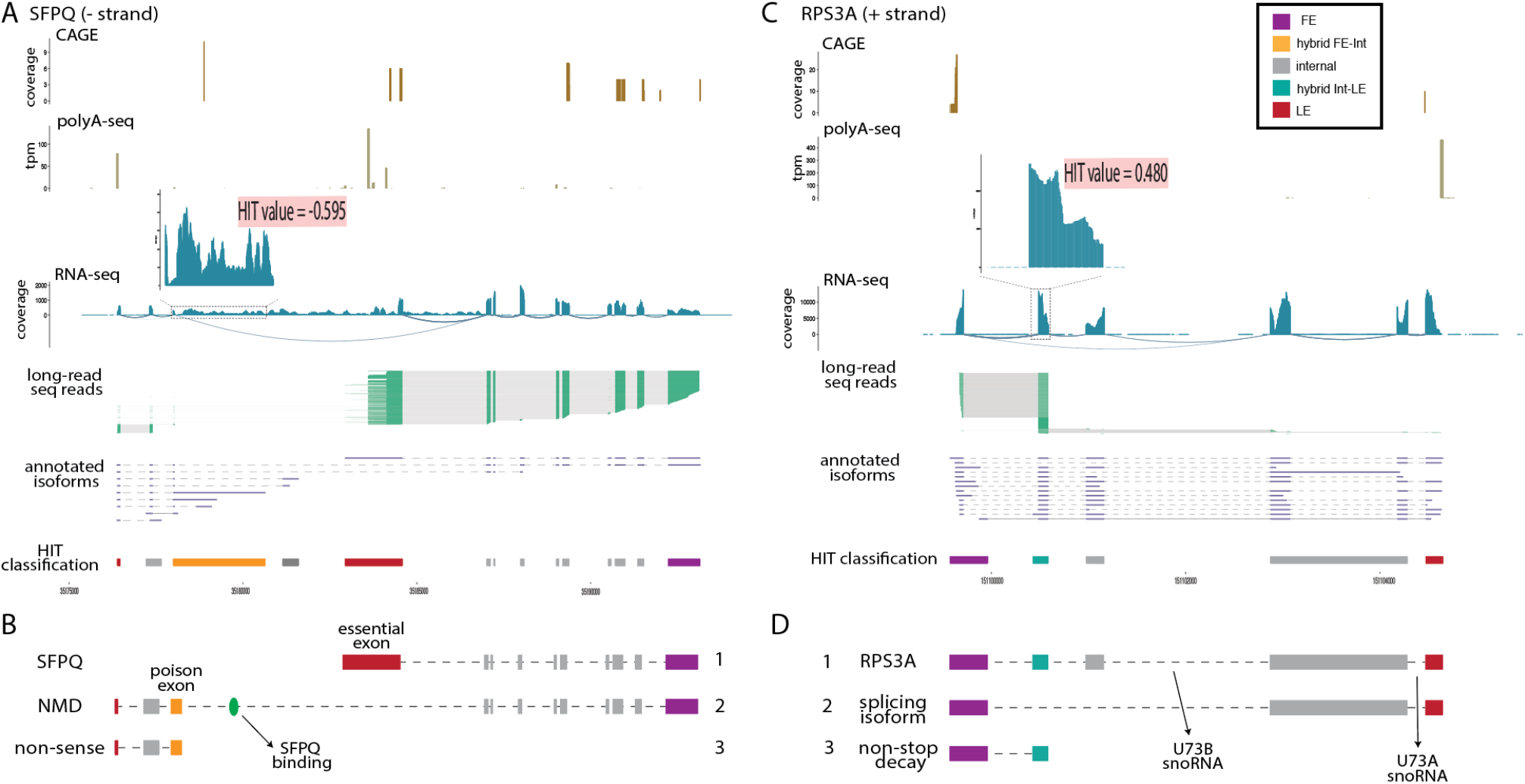
Representative examples of hybrid exons. CAGE-seq peaks, polyA-seq peaks, RNA-seq coverage and junction reads, long-read sequencing reads, annotated isoforms and HIT index classification for SFPQ (**A**) and RPS3A (**C**). (**B)**, SFPQ, encoded by isoform 1 of *SFPQ*, binds its pre-mRNA co-transcriptionally and switches splicing towards the poison hybrid FE-internal exon. (**D)**, Two alternative variants of *RPS3A*, where isoform 3 uses a hybrid LE-internal as a terminal exon and leads to non-stop decay.

In another example, PTPN6, a member of the protein tyrosine phosphatase (PTP) family that functions as an important regulator of multiple signaling pathways in hematopoietic cells, also has a hybrid FE-Internal and several splicing isoforms. We identified three alternative FEs, including a hybrid FE-internal second exon that is associated with expression of the predominant isoform (Figure S6A). Third, we found an unannotated hybrid internal-LE that acts as the predominant LE in LCLs for *RPS3A* (Figure 4C), whose full-length isoform encodes a component of the 40S ribosome subunit. When the hybrid exon is used as an LE, the mRNA lacks a stop codon and is subjected to non-stop decay (Figure 4D). Thus, regulation of the hybrid exon of *RPS3A* controls the levels of the full-length protein of a ribosomal protein. Finally, *CCT8* encodes the theta subunit of the CCT chaperonin, which is involved in the transport and assembly of newly synthesized proteins. In LCLs, we observed expression of two alternative FE and a hybrid internal-LE that produces a truncated protein (Figure S6B). These examples illustrate how hybrid exons can play an important role in modulating gene expression and isoform prevalence of individual genes that have global impacts on gene regulation and protein levels.

### Hybrid exons occur downstream of CpG islands and upstream of polyA sites

To understand the distribution of hybrid exons across human genes, we analyzed the molecular features of the genes in which they occur. We first focused on the gene architecture of these genes, specifically, the number of overall exons and internal exons. We observed that hybrid exons tend to occur in genes with ten or more internal exons (Figure 5A, S7A) and most genes host only one hybrid exon (Figure S7B). However, genes with more internal exons do not have more hybrid exons (Figure S7A), suggesting that additional features govern the regulation of hybrid exons. As expected, hybrid FE-internal exons occur towards the 5’ end of genes compared to hybrid internal-LE exons, although hybrid exons are generally located in internal positions relative to terminal-only exons (Figure 5B, S7C). Furthermore, hybrid FE-internal exons often occur downstream of FEs and upstream of internal exons, while hybrid internal-LE exons often occur downstream of internal exons and upstream of LEs (Figure S7D, S7E). Introns that flank hybrid exons tend to have intermediate length, such that hybrid exons have longer flanking introns compared to internal exons, but shorter flanking introns relative to terminal exons. This may allow for more genomic space for proximal sequence elements that regulate the usage of hybrid exons (Figure 5C, 5D). Hybrid exons also tend to have equally strong or stronger splice sites than terminal exons, reinforcing the idea that the genomic characteristics of hybrid exons fall in between internal and terminal exons (Figure 5E, 5F).

**Fig. 5.**
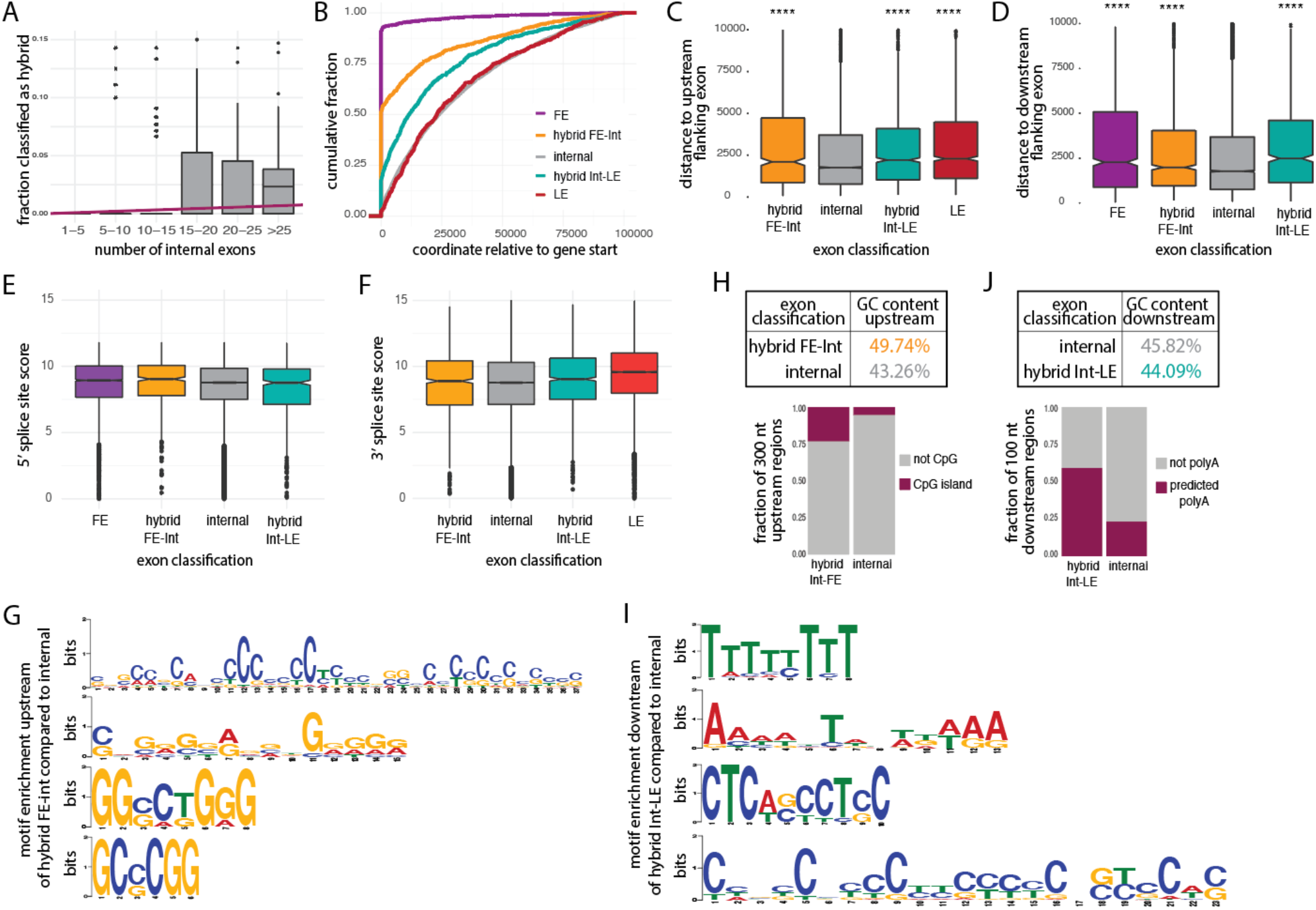
Genomic properties of hybrid exons. (**A)**, Distribution of the fraction of exons classified as hybrid per gene (*y-axis*) binned by the number of internal exons in the same gene (*x-axis*) and linear regression trend. (**B)**, Cumulative distribution function (CDF) of distances to the start of the gene (x-axis) for each of the 5 exons-types across genes expressed in LCLs. Gene start is defined as the start coordinate of the most upstream first exon. (**C)**, Distance from the end of the upstream flanking exon to the start of each metaexon (*y-axis*), binned by exon type (*x-axis*). (**D)**, Same as (C) but distance from the end of each metaexon to the start of the downstream flanking exon. 5’ splice site (**E**) and 3’ splice site (**F**) maxEnt scores (*y-axis*) for metaexons across exon types in LCL (*x-axis*). (**G)**, Enriched motifs in the 100 nucleotide (nt) region upstream of hybrid FE-internal relative to all HIT index classified internal exons. (**H)**, GC content (*y-axis*) in 100nt region upstream of hybrid FE-internal or HIT index internal exons and fraction of 300nt regions upstream of each exon that include a CpG island (*table*). (**I)**, Enriched motifs in the 100nt region downstream of hybrid internal-LE relative to HIT index internal exons. (**J)**, GC content (*y-axis*) in the 100nt region downstream of hybrid internal-LE or HIT index internal exons and fraction of 100nt regions downstream of each exon that include a predicted polyA site (*table*).

We sought to identify candidate genomic features involved in the regulation of hybrid exons, by looking for motifs enriched in regions flanking hybrid exons relative to regions flanking internal exons. For transcription to initiate from a hybrid FE-internal TSS, the immediately upstream sequence should act as a promoter. A recent study showed that ∼60% of random sequences are 1nt away from being able to serve as active promoters in bacteria (32). In mammals, core promoters are still not fully understood, but include structurally and functionally diverse sequences composed of a variety of DNA elements, including CpG islands (33–35). We observed that the regions upstream hybrid FE-internal exons are enriched for motifs with long stretches of Cs and Gs (Figure 5G), with an observed-to-expected CpG ratio greater than 60%, significantly more CpG islands (at least 200 nt of CGs enrichment), and an average of ∼50% GC content (Figure 5H). These CpG islands tend to be located immediately upstream of hybrid FE-internal start coordinates (Figure S7F). Although our results do not show causality, they suggest that CpG rich regions upstream of internal exons are associated with a gain of hybrid usage and novel transcription initiation sites. Conversely, sequences downstream hybrid internal-LE exons are enriched in motifs rich in As and Ts (Figure 5I), have slightly lower CG dinucleotide content, and have significantly more predicted polyA sites than internal exons (Figure 5J). Our findings show that the gain of hybrid internal-LE usage is associated with the presence of downstream polyA sites.

### Widespread usage of intra-tissue and inter-tissue hybrid exons

Alternative mRNA isoforms are often seen as signatures of different tissues or cell types. Thus, we wanted to evaluate the differential usage of hybrid exons across human tissues. To do so, we used our HIT index pipeline to analyze RNA-seq data from GTEx (36). We observed that the distribution of HIT index does not vary significantly across human tissues (Figure 6A), but the numbers of high-confidence hybrid exons show a tissue-specific pattern. Both types of hybrid exons are more predominant in testes, colon and brain tissues, all tissues that are known to have extensive alternative isoform usage (Figure 6B). The brain enrichment is consistent with neuronal genes being larger, with longer introns and undergoing more alternative RNA processing. While both types of hybrid exons showed similar tissue-specificity, there are significantly more hybrid FE-internal exons than hybrid internal-LE exons per tissue (Figure 6B), with twice as many observed in brain tissues, suggesting that hybrid FE-internal are more prevalent in the human genome. The greater variability of hybrid FE-internal exons across tissues suggests that these exons may be more tissue-specifically regulated than hybrid internal-LE exons.

**Figure 6.**
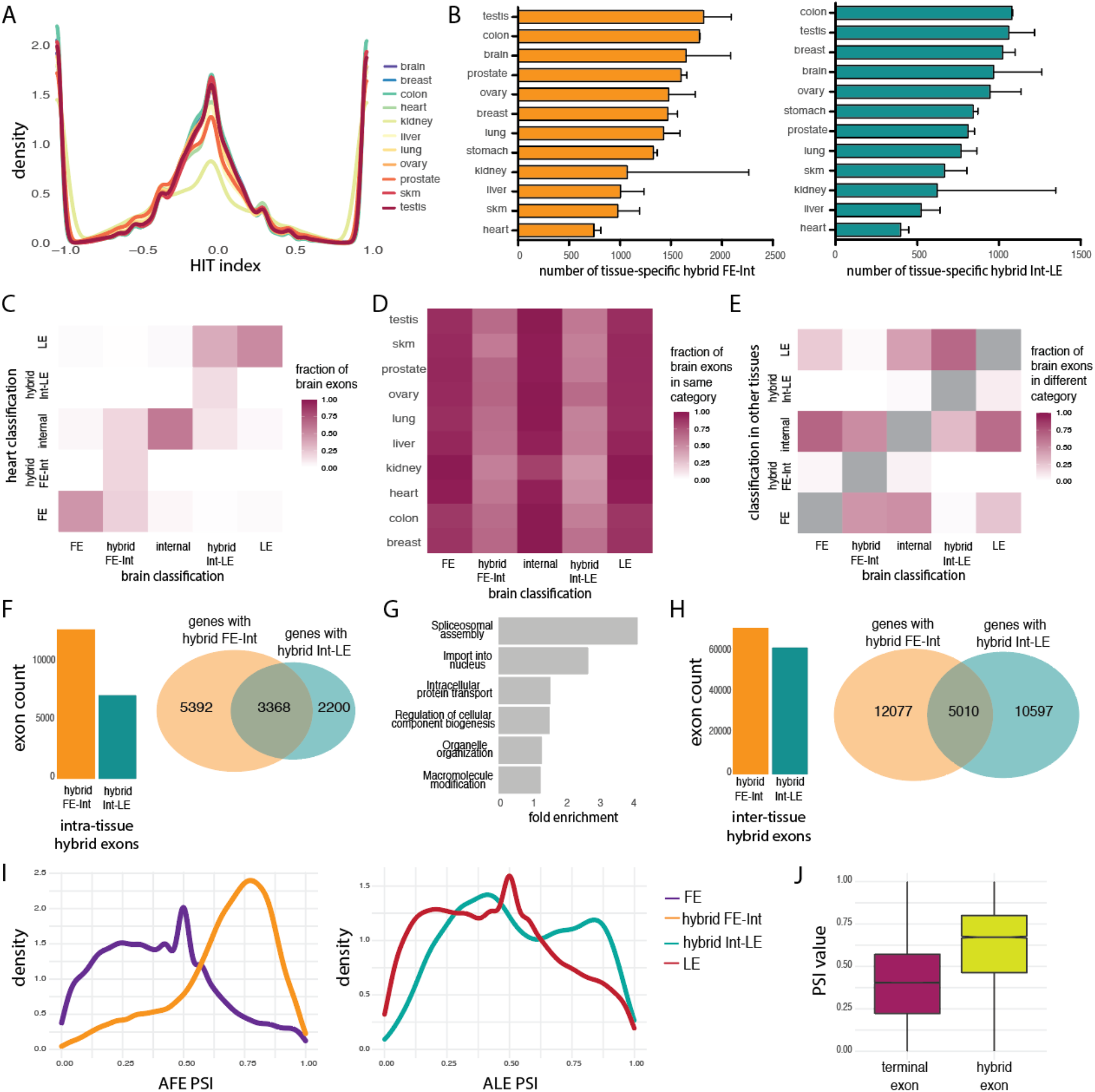
Landscape of hybrid exons across human tissues. (**A**), HIT index distributions of exons expressed (> 10 junction reads) across 11 human tissues. (**B**), The number of tissue-specific hybrid FE-internal (*left*) and hybrid internal-LE exons (*right*) classified by the HIT index across 11 human tissues. (**C**), Heatmap of the fraction of exon-type classifications for exons in brain tissues (*x-axis*) classified as the same or different category in heart tissues (*y-axis*). (**D**), Heatmap of the fraction of classifications of exons in brain (*x-axis*) that are classified in the same category in the other 10 human tissues analyzed (*y-axis*). (**E**), Heatmap of the fraction of classifications of exons in brain (*x-axis*) that are classified in any of the 4 other categories across the other 10 human tissues analyzed together (*y-axis*). (**F**), The number of hybrid FE-internal and hybrid internal-LE exons used in both categories (internal and terminal exon) within the same tissue (*left*) and the overlap between genes with hybrid FE-internal and genes with hybrid internal-LE exons within the same tissue (*right*). The overlap is 1.7 above background expectation (p < 1.6×10^−20^). (**G**), Fold enrichments (x-axis) for gene ontology categories that are significantly enriched (adjusted *p*-values < 0.05) among the 3368 human genes with both types of intra-tissue hybrid exons (y-axis). (**H**), The number of internal exons that are classified as FE or LE in any of the other 10 tissues analyzed (*left*); The overlap between genes with inter-tissue hybrid FE-internal and genes with inter-tissue hybrid internal-LE (*right*). The overlap is 1.3 above background expectation (p < 1.6×10^−20^). (**I**), Distribution of PSI values for alternative FE (*left*) and alternative LE exons (*right*), including hybrid exons across human tissues with multiple AFEs and ALEs. (**J**), Distribution of PSI values (*y-axis*) of hybrid exons relative to terminal exons in genes with multiple FEs and/or LEs across all human tissues analyzed.

Hybrid exons might be more likely to arise in genes with particular functions or tissue-specific expression patterns. To understand whether the same genes consistently utilize hybrid exons across tissues, we calculated the fraction of exons expressed in brain tissues that are classified as the same exon-type in heart or have alternative usage. We observed that, although the majority of exons expressed in brain tissues are not expressed in heart tissues, most terminal or internal exons are classified as the same exon type in heart. In contrast, exons identified as hybrid FE-internal in brain are equally likely to be classified as FE, hybrid FE-internal and internal in heart. Similarly, hybrid internal-LE exons in brain tissues are mostly classified as LE, followed by hybrid internal-LE and internal in heart (Figure 6C). Expanding this analysis to all pairwise tissue comparisons, both types of hybrid exons are consistently less likely to be classified as hybrid across tissues relative to terminal or internal exons, which suggests that hybrid exons are more likely to be alternatively used across tissues and hybrid usage tends to be tissue-specific (Figure 6D). We note that since hybrid exons are more difficult to detect, the detection of hybrid exons may be underestimated. Exons classified as FEs and LEs in brain tissues are both mostly classified as internal in other tissues, while brain internal exons are mostly classified as FE and LEs. Similarly, brain-specific hybrid exons are mostly classified as internal and FE or LE (depending on their hybrid type; Figure 6E). Finally, the HIT index classifications of exons in the same tissue across different individuals in the GTEX dataset are significantly more similar than the exon classifications of a given individual across tissues (Figure S8A, B, C), suggesting that exon classifications are highly tissue-specific and robust to technical variance across libraries. Together, our findings demonstrate that hybrid exons are more likely to be used as both terminal and internal exons in a tissue-specific manner, while being used as only terminal or internal exons in other tissues. This is consistent with the observations that hybrid exons tend to have properties intermediate between internal and terminal exons, and potentially allowing for more flexibility in isoform usage.

Since hybrid exons allow for increased isoform diversity, we reasoned that hybrid exons might arise within genes that would benefit from flexibility in RNA processing. We first asked if genes with one type of hybrid exon are more likely to have another type of hybrid in the same tissue. Indeed, genes with hybrid FE-internal exons are 1.3-fold more likely to have hybrid internal-LE exons than expected by chance (overlap p-value < 9.822×10^−85^) (Figure 6F). Genes with both types of intra-tissue hybrid exons are enriched in spliceosome components and splicing factors (Figure 6G). The enrichment of hybrid exons within factors involved in splicing may contribute to the autoregulation of these factors, as for SFPQ above, or may contribute to tissue-specific splicing programs. More broadly, an exon could be used as a terminal exon in one tissue, but as an internal exon in another tissue, which we term inter-tissue hybrid exons We identified more than 130,000 inter-tissue hybrid exons across human tissues (10% of all exons analyzed across tissues) and similar to intra-tissue hybrids, genes with inter-tissue hybrid FE-internal are significantly more likely to have an inter-tissue hybrid internal-LE (Figure 6H). Finally, we asked what proportion of isoforms are likely to use hybrid exons as terminal exons. Surprisingly, we observed that hybrid exons are used in a relatively high proportion of isoforms compared to terminal exons, as measured by percent spliced in (PSI) values. Hybrid exons produce the predominant gene isoforms across all tissues analyzed (Figure 6I & J S8D & E). Our findings indicate that tissue-specific hybrid exons drive large changes in isoform usage and are likely to have major impacts on mRNA and protein synthesis.

## Discussion

Here, we describe the first computational pipeline to identify and classify hybrid exons, a largely undescribed type of exons, from RNA-seq data. The HIT index uses splice junction reads to classify exons as terminal exons that serve solely as FEs or LEs in a transcript, internal exons, or hybrid exons. We find that, conservatively, 10% of human exons are intra-tissue hybrid exons that can serve as either terminal or internal exons in a transcript. However, technical biases due to library preparation and low proportional usage of exons (Fig. X, simulation data) may lead to an underestimation of the true number of hybrid exons in human tissues. Moreover, increased tissue and cell type sampling with higher coverage sequencing data could increase our power to identify intra- and inter-tissue hybrid exons and the contexts in which they are used.

Orthogonal genome-wide high-throughput sequencing datasets targeting terminal site usage and long-read RNA sequencing were used to validate our classifications and hybrid exons in individual genes. We obtained 100% TSS and polyA specificity for all terminal exons including hybrid exons (Figure 3). Consistent with previous studies (37–40), we find that the high frequency of truncated reads in the long-read sequencing prevented us from using these data to globally validate terminal or internal site usage. Combined with high error rates and potential coverage biases, long-read Direct RNA sequencing anchored on the polyA tail leads to substantial 5’ truncation of transcripts, reflected in 5’ ends of reads being located farther, on average, from annotated terminal sites than 3’ ends of reads and reads lengths that are smaller than average transcript lengths of the corresponding annotated isoforms (Figure S9A, B). These biases significantly affect terminal exon identification and quantification. Thus, only 50% of our HIT index classifications are matched using long-read sequencing methods (Figure S9C), which could be due to either the aforementioned lack of power in long-reads for terminal exon identification, limitations of the HIT index classifications, or a combination of these effects. Regardless, despite the great promise of long-read sequencing technologies, short-read sequencing datasets are still necessary to classify exons. Furthermore, the HIT index may provide a useful tool to calibrate terminal end truncation in long-read RNA sequencing datasets.

We found that hybrid exons are an inherently flexible exon type due to their ability to serve as either terminal or internal exons within a transcript. Given their potential to increase both the coding capacity and regulatory potential within a genome, hybrid exons are an underappreciated way in which cells can expand their transcriptome and proteome to fine-tune alternative isoform usage from a single gene. Since hybrid exons can influence either untranslated regions or coding sequences depending on their usage, they are likely subjected to different evolutionary selective pressures than either internal-only or terminal-only exons. We found that both FE and hybrid FE-internal exons have fewer ATGs that start the open reading frame (ORF) compared to internal exons (Figure S9D), suggesting that they mostly affect untranslated regions. However, hybrid FE-internal exons have higher frequency of upstream ORFs (uORFs), which are key regulators of gene expression and often activate or inhibit downstream expression of the primary ORF. Their dual-regulation by both splicing machinery and transcription or polyadenylation machinery (for hybrid FE or LE exons, respectively), suggests that hybrid exons lie in regions with flexible or multi-purpose regulatory sequences.

Our analysis detected CG-rich sequences upstream of hybrid FE-internal and polyA rich motifs downstream of hybrid internal-LE exons, indicating that hybrid exons are enriched for specific sequence features that may enable their use as either terminal or internal exons. These features are generally associated with transcription initiation and polyadenylation, respectively, but we did not specifically identify any auxiliary sequences canonically associated with mRNA splicing (apart from the expected canonical splice sites). These findings suggest that there might be competition sites involved in splicing and transcription/polyadenylation for usage of a particular exon, which may be influenced by as yet unidentified sequence features, local epigenetic marks, DNA confirmation, RNA secondary structure, or coordination with other RNA processing events on the same transcript. We speculate that instead of creating new exons (8, 41), hybrid exons represent exons that gained new functions in order to increase diversity over evolution. Further studies of the evolutionary histories of these flexible hybrid exons may shed light on how new exons or isoforms arise in tissues or species and the mechanisms by which they are recognized by transcription factors, splicing components and termination machinery.

Together, our findings demonstrate the unique role that hybrid exons play in human transcriptomes and highlight that our understanding of transcriptome complexity is far from complete.

## Materials and Methods

### HIT-index pipeline for exon classification

The HIT index approach uses the ratio between splice junction reads that overlap the beginning or end of exons to classify exons as being first, hybrid first, internal, hybrid last, or last exons. The HIT index pipeline involves 3 primary steps: (1) annotating meta-exons from a transcript annotation, (2) extracting overlapping junction reads for each exon, and (3) calculating the HIT index, classifying exons, and estimating percent spliced in values for alternative terminal exons. The full pipeline is available as a set of python scripts, with detailed usage instructions, at github.com/thepailab/HITindex.

### Exon definition and junction read counting

First, for each gene in a transcriptome reference file (i.e. gtf), annotated exons with overlapping boundaries are merged with pybedtools (42) to create a meta-exon. The boundaries of these meta-exons are used to extract junction reads. To account for ambiguity in the exact position of transcription start sites, splice sites, or mRNA cleavage sites, an optional buffer region is added to the boundaries of meta-exons, with a recommended distance of 50nt upstream of the 5’end of the exon and 30nt downstream of the 3’end of the exon. Second, splice junction reads (SJRs) (split between two regions) are extracted using samtools (43). For each exon, SJRs that span the 5’ boundary (with at least 10nt overlap) are counted as upstream SJRs and SJRs that span the 3’ boundary (with at least 10nt overlap) are counted as downstream SJRs, regardless of the junction-specific boundaries (ie. specific splice sites used). For example, an exon with 2 alternatively used 5’ splice sites would have an upstream SJR count that is the sum of junction reads derived from splicing to both 5’ splice sites. SJRs that begin and end in the same meta-exon are considered spurious and not used for further analysis.

### HIT index estimation

The HIT index and associated metrics are calculated for each exon with at least 5 total SJRs. Specifically, the HIT index is calculated as the the ratio of the imbalance in upstream and downstream SJRs over the total SJRs for the exon, as defined in Equation 1.

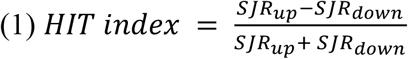

Two metrics are used to assess robustness of the HIT index estimation: (1) bootstrapped confidence intervals and (2) a bootstrapped variance metric representing the probability that the true value of the HIT index as within ±10% of the observed value. Both metrics are calculated through bootstrapping HIT indices for each exon from reads randomly subsampled with replacement for a user defined number of bootstraps (1000 suggested samples). 75%, 90%, and 95% confidence intervals are calculated from the distribution of bootstrap statistics. The bootstrapped variance metric is calculated as the number of times that bootstrapped statistics are at least 10% lower or higher than the actual HIT index estimate.

### Edge-effect flag

Simulated data uncovered biases in the HIT index estimation that existed specifically for exons near the edges of transcripts, where an RNA-seq edge effect led to a depletion of upstream or downstream SJRs for exons near the 5’ or 3’ ends of transcripts, respectively (Figure S1C). To account for this, the HIT index pipeline flags exons that have a higher probability of being affected by the edge effect and allows the user to decide whether to include them in downstream analyses. Exons are flagged based on their distance to the transcript end and an empirical determination of the distance at which SJR ratios are biased due to the edge effect, which is expected to be approximately one fragment length away from the end of the transcript. Specifically, piecewise linear regression is used to find the inflection point between the two subsets of exons: (1) exons unaffected by edge-effect induced SJR biases, with equal upstream and downstream SJRs on average and (2) exons affected by edge-effects, with a linear relationship between the exon’s distance from the transcript end and the SJR bias. Exons with a 5’ or 3’ distance lower than the inflection point and not classified as terminal only exons (|HITindex| = 1) are flagged as edge-effect exons.

### Probabilistic Framework

Given technical biases in the data, we developed a generative approach that estimates the probability that the ratio of downstream SJRs to the total SJRs arises from a particular exon type. Rather than assume that the ratio of downstream SJRs are generated from exons with some fixed probabilities, we assume that first-only, internal-only, and last-only exons generally have downstream SJR probabilities that are near 1, 0.5, and 0, respectively, but vary among the different exons in each class. Assuming the modes of these distributions are likely 1, 0.5, and 0, the model estimates two features from the data: (1) how concentrated downstream SJR probabilities are around these modes and (2) the probabilities with which exons belong to each of these classes. Using these two assumptions, we model downstream SJR probabilities as arising from a mixture of these three distributions. Hybrid exons are then defined as exons poorly explained by or outliers relative to any of the three non-hybrid distributions.

Specifically, we define *D*_*i*_ as the number of downstream SJRs mapping to exon *i* out of *N*_*i*_ total SJRs. We model *D*_*i*_ as arising from a mixture of beta-binomial distributions, representing the classes of first-only *F*, internal-only *I*, last-only *L*, and hybrid *H* exons, as defined in equation 2.

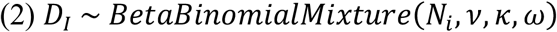

A Dirichlet prior is used for the probability, *ω*, that an exon belongs to each class, defined as *ω* ∼ *Dirichlet*(*α*). The modes of the beta distributions in the beta-binomial mixture are defined as, which are held constant as fixed assumptions: *ν* = {*ν*_*F*_ = 1, *ν*_*I*_ = 0.5, *ν*_*L*_ = 0, *ν*_*H*_ = 0}. The concentration of beta-distributions around these modes *κ*, which we want to learn from the data for the first-only, internal-only, and last-exons. We place relatively uninformative priors on these parameters, drawing them from half-normal distributions shifted to start at 3 since *κ* must be greater than 2 in order for the beta-distributions for *κ*_*F*_ and *κ*_*L*_ to not become uniform: *κ*_*F*_, *κ*_*I*_, *κ*_*L*_ ∼ *HalfNormal*(*loc* = 3, *σ* = 1000). In contrast, for the class of hybrid exons, we set *κ*_*H*_ = 2, which, when coupled with the model *ν*_*H*_ = 0, corresponds to a uniform distribution under the assumption that for hybrid exons, all downstream SJR probabilities are equally plausible. We use Automatic Differentiation Variational Inference (ADVI) as implemented in the PyMC3 python package (44) to fit an approximation of the posterior distribution, allowing estimates of the posterior means and variation around this mean. Given a sample *θ* of *M* draws from the posterior distribution, defined as *θ*_*i*_ ∼ *P*(*θ* | *D*_*i*_*N*_*i*_), we can estimate the probability that an exon arose from a particular component distribution *z*_*i*_ based on observed *D*_*i*_ and *N*_*i*_ counts, as defined in equation 3.

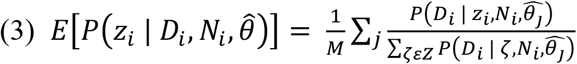

Finally, hybrid exons are partitioned into two classes using the posterior over the underlying proportion of downstream SJRs, with *P(HybridFI)* and *P(HybridIL)* defined in equations 4 and 5, respectively.

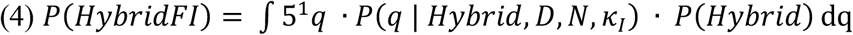

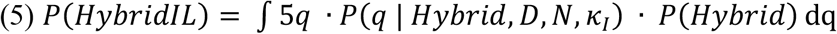

The generative model leads to the following outputs: (1) the posterior mean estimate for the fraction of downstream SJRs, (2) the posterior mean estimate for the fraction of downstream SJRs converted to the HIT index, (3) the posterior classification probabilities for first-only *P(F)*, first-internal *P(FI)*, internal *P(I)*, internal-last *P(IL)*, and last-only *P(L)* exons.

### Exon Classification and PSI value calculations

Exons are classified into the following categories: “first”, “FirstInternal_medium”, “FirstInternal_high”, “internal”, “InternalLast_medium”, “InternalLast_high”, and “Last” exons, using a combination of thresholds across the HIT index metic, statistical confidence metrics, and posterior probabilities from the generative model, with all thresholds specified in the user defined HIT_identity_parameters file. Specifically, terminal exons are defined as having a |HITindex| greater than or equal to the specified “HITterminal” threshold and a bootstrapping variance value less than the specified “HITpval”. Hybrid exons are defined as having a |HITindex| greater than or equal to the specified “HIThybrid” threshold, a |HITindex| less than the specified “HITterminal” threshold, and an appropriate generative model posterior probability greater than the specified “prob_med” or “prob_high” thresholds, for medium and high confidence classifications, respectively. Finally, internal exons are defined as all remaining exons, including those exons whose specified bootstrapped confidence interval overlaps zero. Exons classified as “first”, “FirstInternal_medium” or “FirstInternal_high” are used to calculate percent spliced in (PSI) values for alternative first exon (AFE) usage, while exons classified as “last”, “InternalLast_medium” or “InternalLast_high” are used to calculate PSI values for alternative last exon (ALE) usage, for each exon *i* out of *n* total first or last exons as defined in equations 6 and 7, respectively.

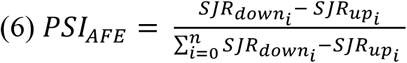

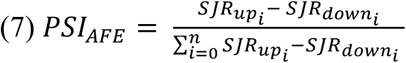

If the user selects the --edge flag, exons flagged as potentially being affected by edge-effects (see above) are not included in the PSI value calculations.

### Simulations

We simulated 12 sets of artificial transcripts to assess the specificity and sensitivity of both the FILindex and generative model (Figure S1A). For each of these transcript architectures, we simulated short read RNA-sequencing data from a range of different genomic and expression contexts: exon length (100nt - 1kb), proportional usage of alternative first or last exons (0.05 - 0.95 PSI), proportional usage of skipped exons (0.05 - 0.95 PSI), library insert size (150 - 300nt), read length (50 - 100nt), and sequencing coverage (10-100M reads). Transcripts were also sampled from a range of gene expression levels that matched the distribution of TPMs in real RNA-seq data (Figure S1B). To generate short read data from transcripts, we simulated several steps of Illumina library generation, including transcript fragmentation (using a modified Weibull distribution) and fragment size selection (as detailed in (45)). The FILindex and generative model were applied to simulated read data as described above.

### RNA-seq data processing and analyses

We used RNA-seq data from LCL human cells available at NCBI Gene Expression Omnibus (accession no. GSE30400 (46)) and several human tissues from different individuals from GTEx (36) downloaded through dbGaP under accession numbers SRR1068855, SRR1091865, SRR1366790, SRR1093075, SRR1375738, SRR1093527, SRR1437274, SRR1341721, SRR1434586, SRR1347481, SRR1347481, SRR1313449, SRR1080415, SRR1085975, SRR1313969, SRR1322373, SRR1362332, SRR1395999, SRR1441768, SRR1470273, SRR1082352 SRR1086140, SRR1317751, SRR1323087, SRR1366473, SRR1396700, SRR1460409, SRR1490246, SRR1071379, SRR1083632, SRR1090650, SRR1321351, SRR1336314, SRR1388190, SRR1416516, SRR1464788. We excluded cases with different numbers of reads in read1 and read2 and selected the most recent releases in cases with multiple ones. Reads were mapped using STAR (47), guided by transcriptome coordinates in the genome annotation database (Ensembl GRCh37.72). The HIT index pipeline was run on resulting bam files, with default parameters, to classify exons into five categories. Expressed exons required SJR > 0 and classified as follows:

FE: HITindex == -1

Hybrid Internal-FE: -0.3 > HITindex > -0.95 & Probability of being hybrid > 0.5

Internal: -0.3 < HITindex < 0.3

Hybrid Internal-FE: 0.3 < HITindex < 0.95 & Probability of being hybrid > 0.5

LE: HITindex == 1

All gene ontology analyses were performed using the Gene Ontology (GO) knowledgebase developed by the GO Consortium, using relevant expressed genes as a background set.

### CAGE, 3p-seq, and polyA site data processing and analysis

To evaluate performance of the HIT index, start and end coordinates of all classified exons in LCL cells were interrogated for overlap on the same strand with TSS sites from CAGE and polyA sites from 3’-seq within 50, 100, 150, 200, and 250 bp. Specificity was calculated as the fraction of exons overlapping a bonafide TSS or PolyA site within the indicated window size. TSS sites were identified as the nucleotide within a peak with the maximum score in CAGE data from the FANTOM project (26) while polyA sites were also identified with nucleotide resolution in 3’-seq data downloaded from the polyAsite database (27).

### Motif, splice site, context and ORF analyses

DNA sequences flanking classified exon were extracted from the genome annotation database (Ensembl GRCh37.72) using GetFasta function from BEDTools (48) with default parameters and strand-specificity. Motif analyzes were done with MEME suite packages (49) and customized R scripts. The 5’- and 3’-splice site of each exon was mapped to hg38 reference genome using BEDTools (v.2.28.0) and splice site scores were calculated with MaxEntScan (50). Among all splice sites for overlapping exons including in each meta-exon, the exons with the highest splice site score were selected. ORF and uORF analyses were performed in R (v.4.0.5) using the ‘matchPattern’ function from R package Biostrings and customized scripts. All occurrences of start and stop codons were annotated for each exon. All graphical plots were generated using the R package ggplot2.

### Long-read RNA sequencing analysis

A raw fastq file with base-called Oxford Nanopore direct-RNA sequencing data from the GM12878 lymphoblastoid cell line (51) was downloaded and mapped to the GRCh38 reference assembly using minimap2 (52). 98% of the 13.3M total reads mapped with high confidence primary alignments. Mapped reads were assigned to genes using the GRCh38 gtf annotations and for downstream analyses, only reads mapping to protein coding transcripts were considered. To account for technical biases in the long read sequencing data that lead to truncated or chimeric reads, we undertook a number of filtering steps to obtain a high-confidence set of reads. First, we calculated z-scores for reads across the distribution of reads per gene to estimate how many standard deviations a mapped length is away from the mean mapped length of the gene. We discarded reads with mapped lengths 3 z-scores away from the mean. Second, we compared the mapped length of a read against annotated coordinates using GRCh38 annotations. Reads with mapped lengths less than or greater than 1.5kb from the smallest or largest annotated transcripts for the corresponding gene, respectively, were discarded. Finally, we only considered genes with more than 10 reads. After applying these filters, we obtained 7.7M reads across 10.4k genes, with a mean read length of 1.1kb. To identify first, internal and last exons, each read was split into discrete exons using the CIGAR string. For each read, the most upstream and downstream exons were classified as first or last exons, respectively, and all other exons were considered as internal exons.

## Acknowledgments

We thank all members of the Burge lab, Fiszbein lab and Pai lab for useful discussions and comments.

## Author contributions

Conceptualization: AF, AAP

Methodology: AF, MM, CBB, AAP

Investigation: AF, MM, ECR, GK, AAP

Visualization: AF, MM, ECR, GK, CBB, AAP

Supervision: AF, CBB, AAP

Writing—original draft: AF, AAP

Writing—review & editing: MM, ECR, GK, CBB

## Data and materials availability

The HIT index python pipeline is available at github.com/thepailab/HITindex and all R scripts for analysis are available upon request.

## Supplementary Figures

**Fig. S1.**
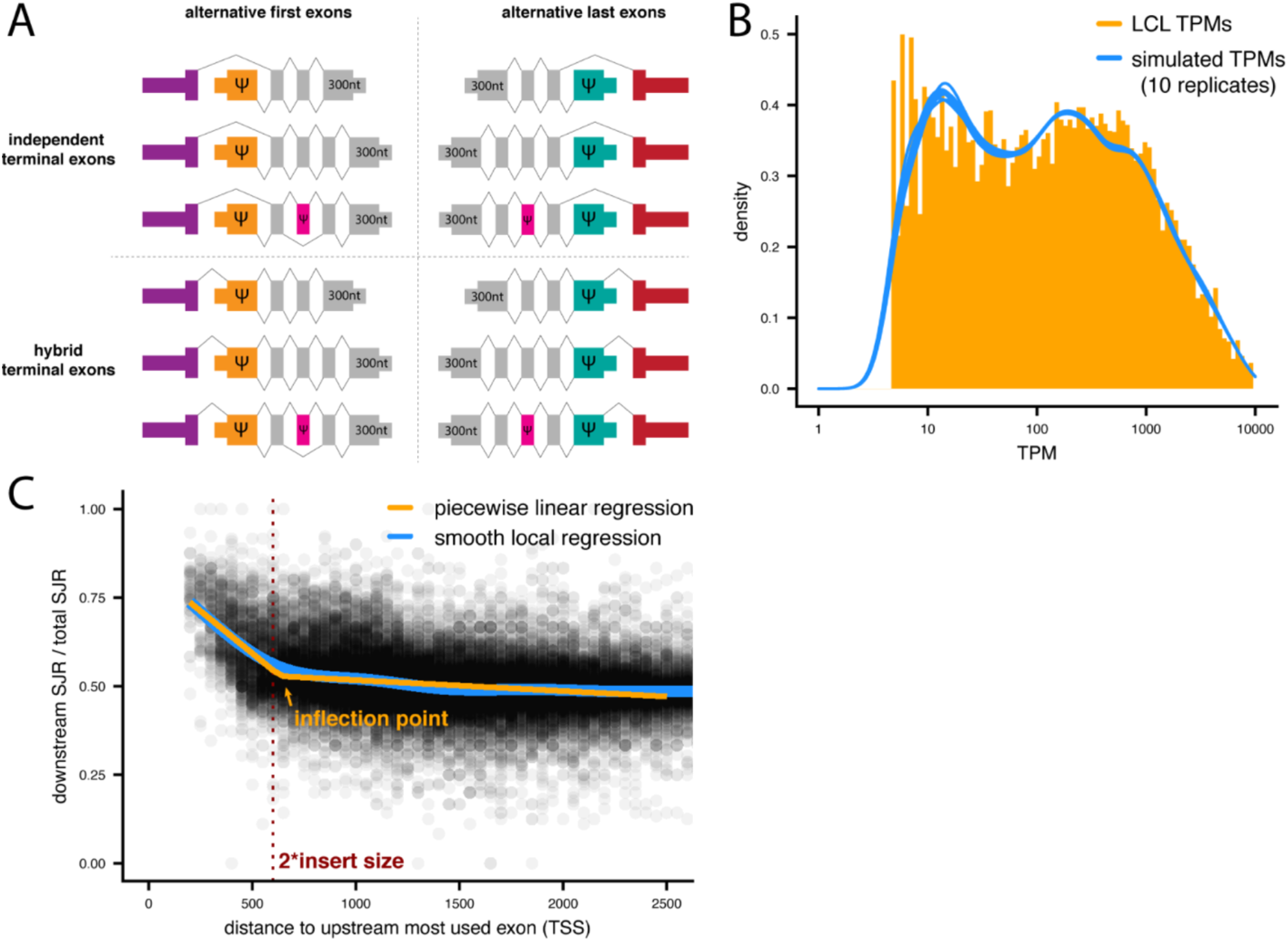
(**A**), Schematics of the 12 sets of artificial transcripts created for simulated RNA-seq data, with constitutive exons denoted in grey. Alternative terminal, hybrid, or internal cassette exons were included in the transcripts with variable PSI values. Transcripts with alternative or hybrid first exons had constitutive 300 nt last exons, and visa versa. (**B**), The distribution of simulated transcript per million (TPM, *blue*) values were drawn to match the distribution of true TPM values in LCLs (*orange*). (**C**), The skew in the ratio of simulated downstream SJRs (*y-axis*) for non-terminal exons relative to the distance of the exon away from the TSS (*x-axis*), where the data shown were simulated 300nt insert size (*dotted red line*), as seen using a local loess regression (*blue line*). The inflection point determined from the data using a piecewise linear regression (*orange line*) falls very close to two insert sizes away from the TSS (*dotted red line*).

**Figure S2.**
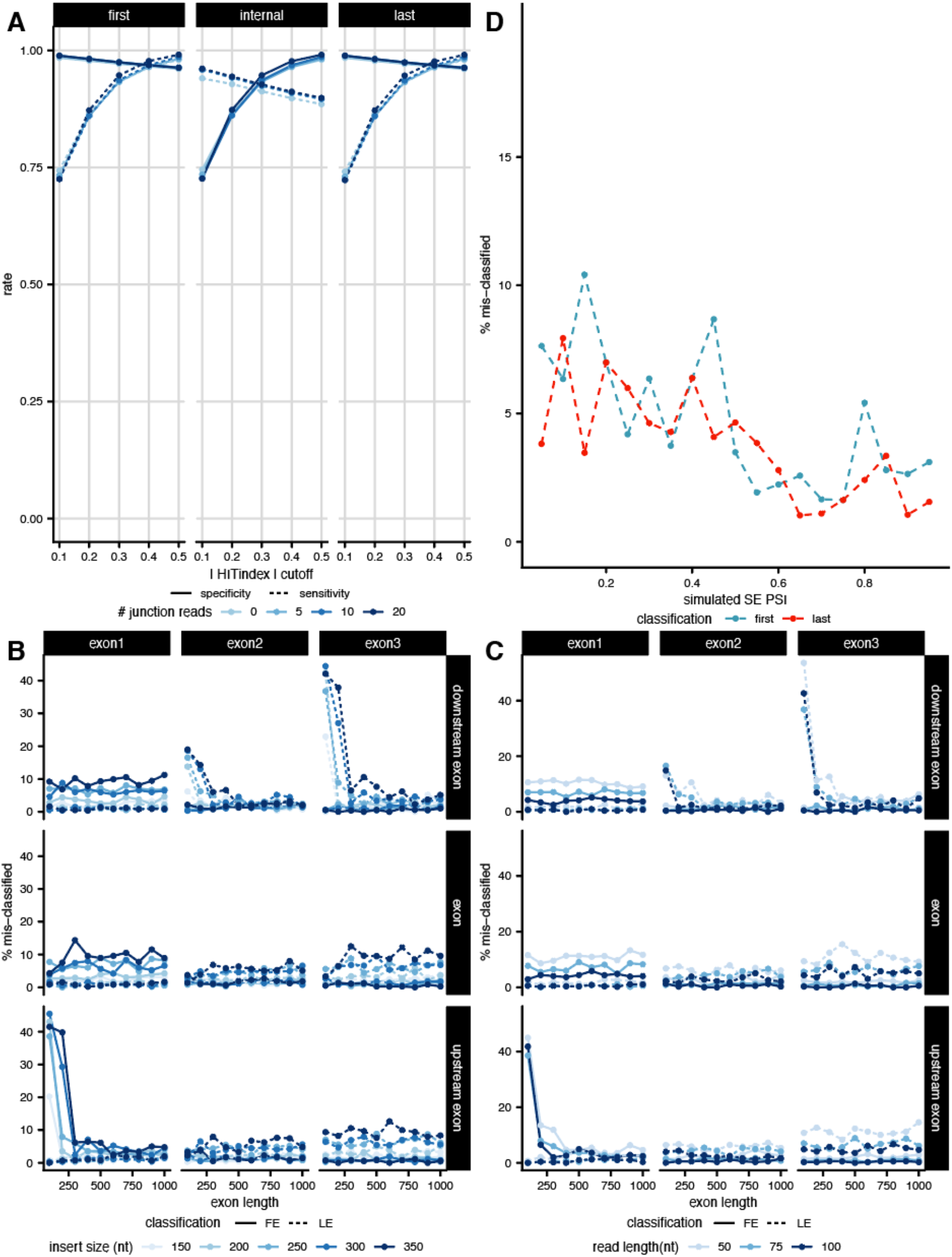
(**A**), True positive (sensitivity, *dotted lines*) and true negative (specificity, *solid lines*) rates (*y-axis*) for simulated exon classification across moderate HIT index thresholds (*x-axis*), for different total SJR thresholds. The percent of internal exons incorrectly classified (*y-axis*) as either first (*solid lines*) or last exons (*dotted lines*) due to edge effects across varying exon lengths (*x-axis*) and insert sizes (**B**) or read lengths (**C**). (**D**), The percent of alternatively used cassette exons incorrectly classified (*y-axis*) as either first (*blue*) or last (*red*) exons across varying PSI values (*x-axis*).

**Figure S3.**
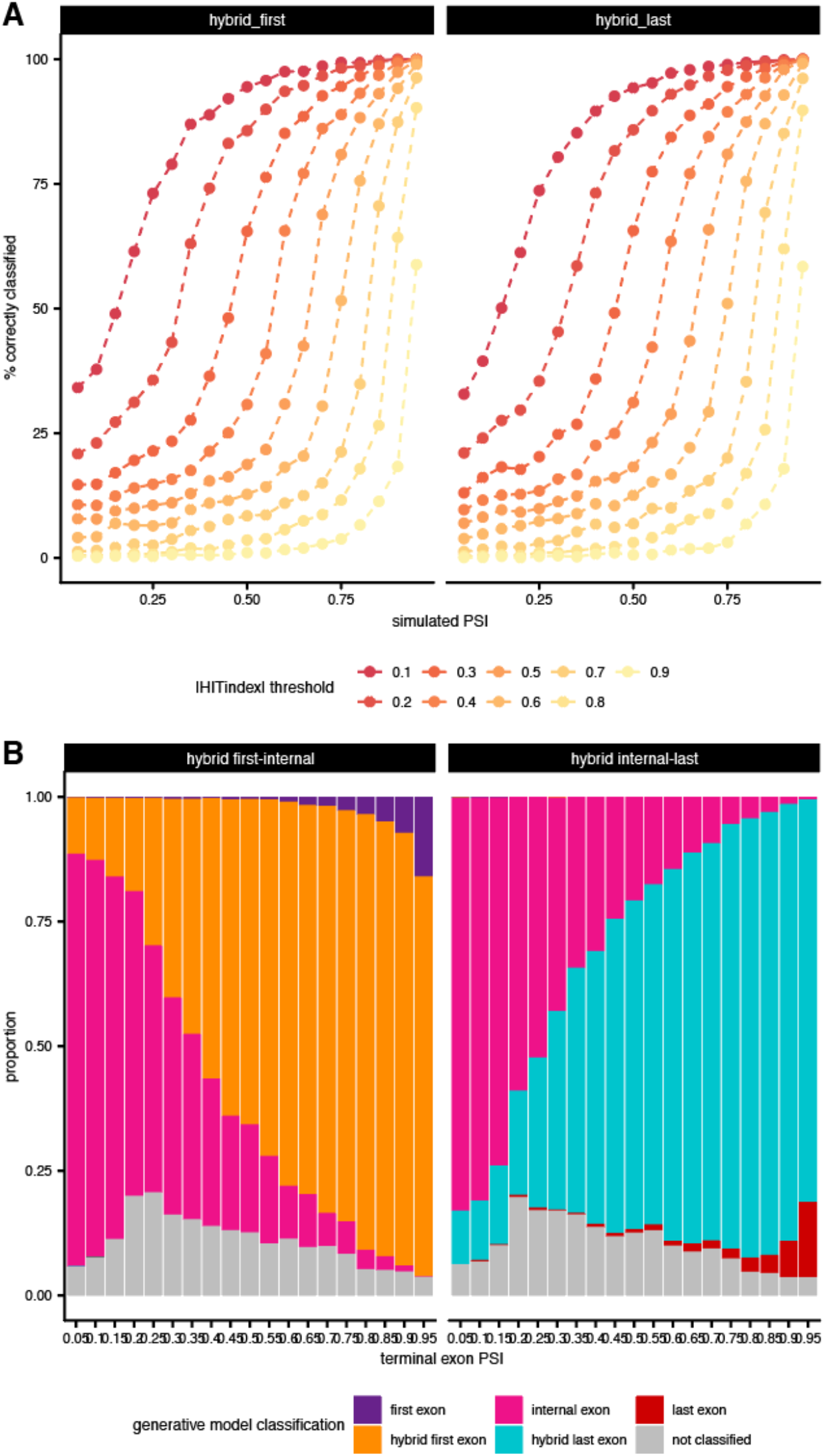
(**A**), The percent of hybrid exons incorrectly classified (*y-axis*) for either hybrid first-internal (*left*) or internal-last (*right*) exons, across varying PSI values (*x-axis*) and |HITindex| thresholds (*colors*). (**B**), The proportional classification of hybrid first-internal (*left*) or internal-last (*right*) exons into each of the 5 exon types by the generative model, as defined by the maximum posterior probability for each exon. Exons are “not classified” if no posterior probability is greater than 0.7.

**Figure S4.**
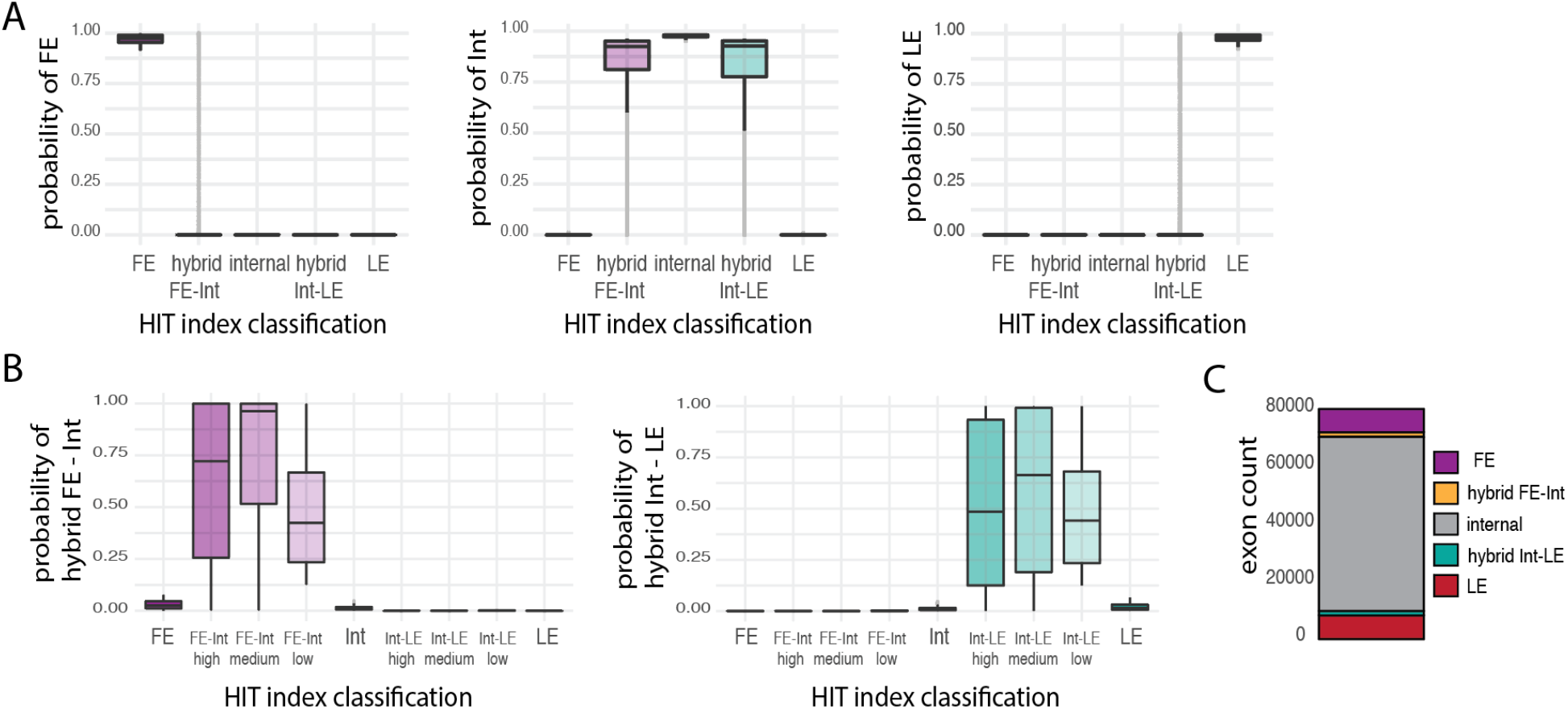
(**A**), probability of being classified as terminal or internal exons by the generative model for all 5 categories of exons analyzed. (**B**), probability of being classified as hybrid exons by the generative model for all 5 categories of exons analyzed including three subsets of hybrid exons with different confidence levels based on their adjusted p-values. (**C**), number of expressed exons classified by the HIT index and the generative model in 5 categories including 2 of hybrid exons.

**Figure S5.**
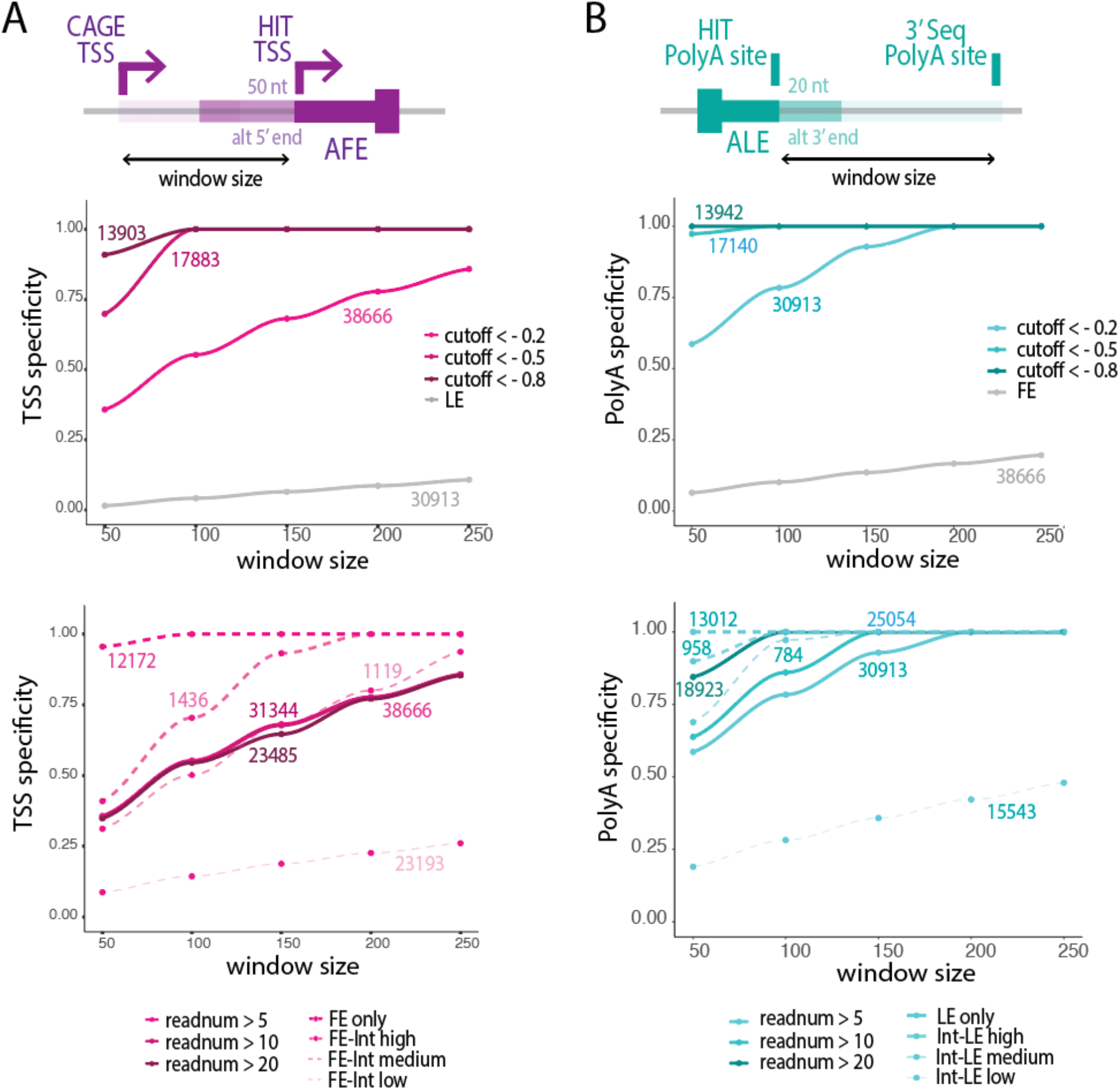
TSS specificity (**A**) and polyA specificity (**B**) for each window size, where specificity is calculated as the fraction of HIT TSSs (start coordinate of identified FE) or HIT polyA (end coordinate of identified LE) that fell within the window size of any TSS identified by CAGE or polyA identified by 3’seq. TSS specificity (**A**) and polyA specificity (**B**) are shown for different cutoffs and LE or FE as a negative control (upper panel), and for different expression thresholds and confidence intervals of hybrid FE-internal or hybrid internal-LE (lower panel).

**Figure S6.**
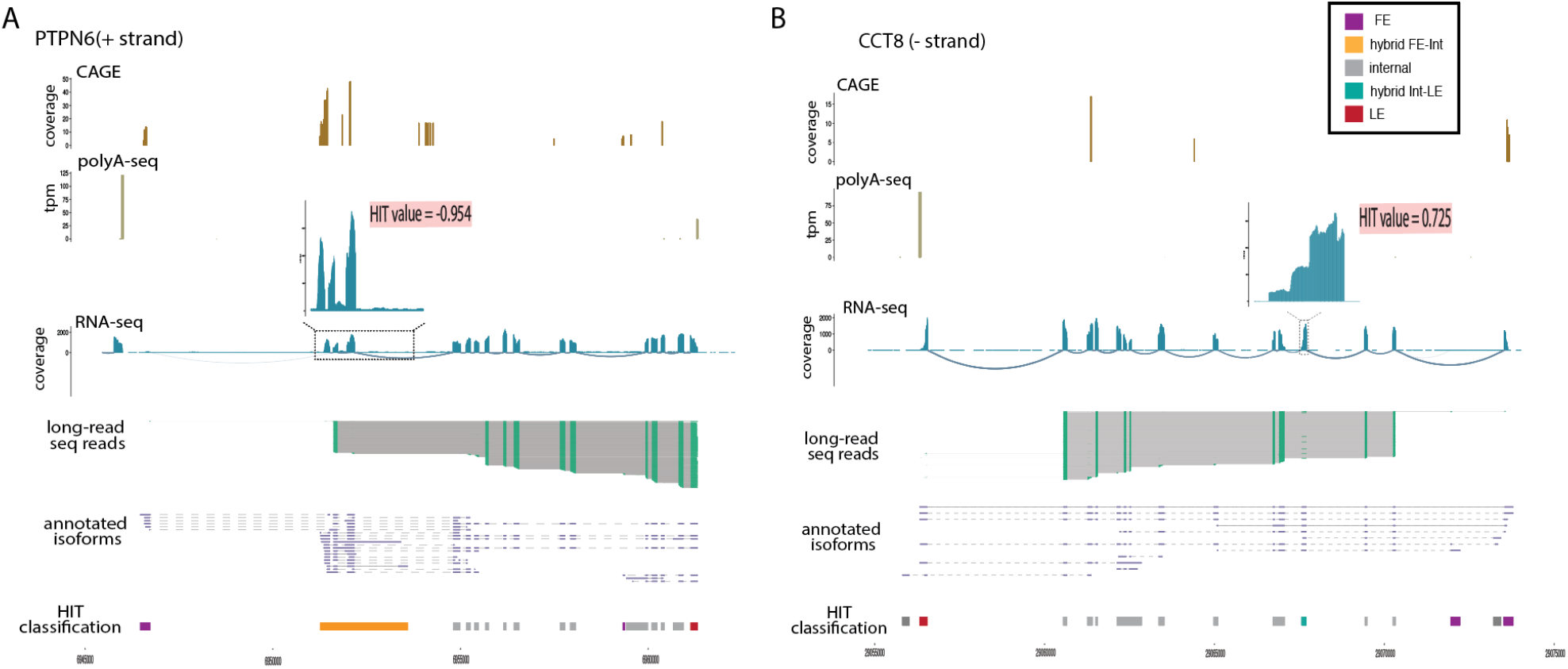
CAGE peaks, polyA-seq peaks, RNA-seq coverage and junction reads, long-read sequencing reads, annotated isoforms and HIT index classification of PTPN6 (**A**) and CCT8 (**B**).

**Figure S7.**
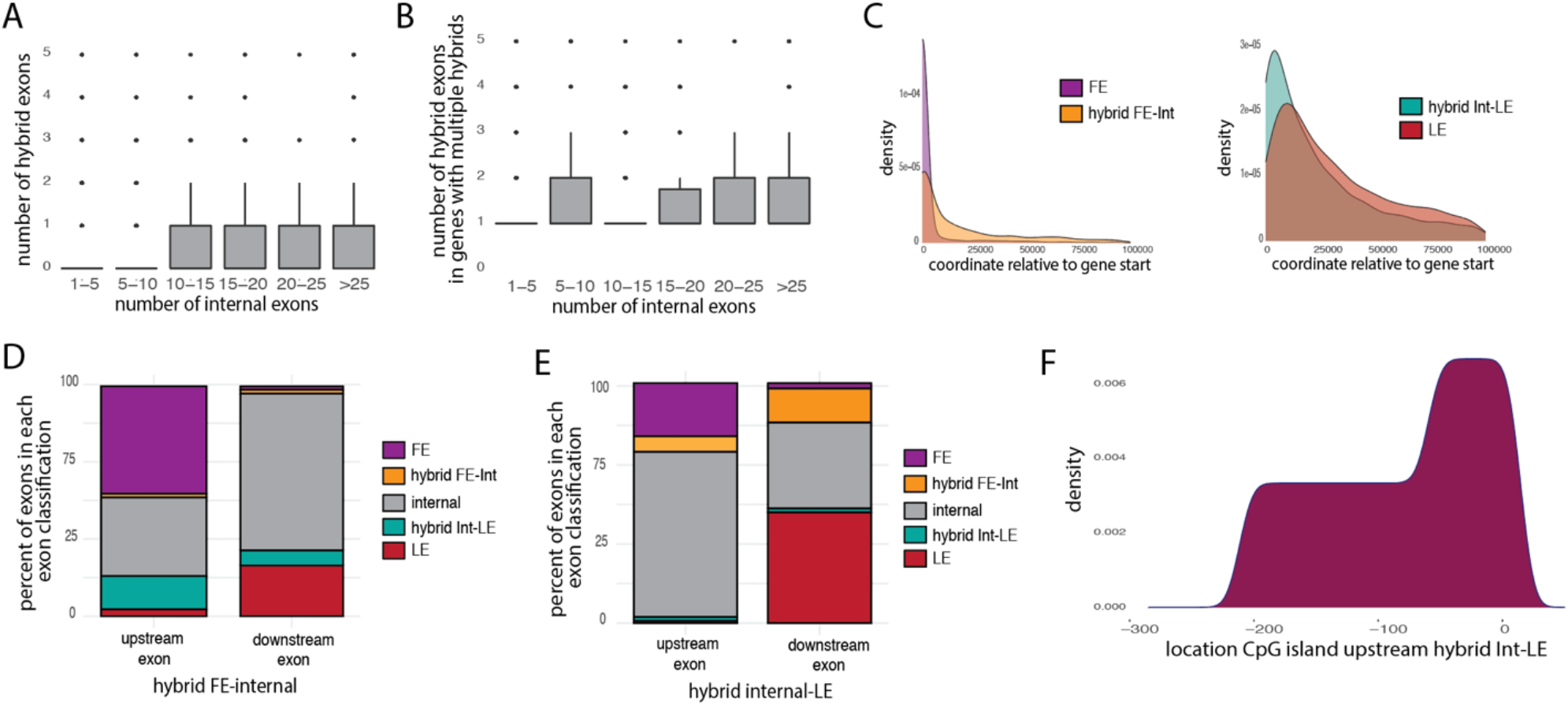
(**A**), distribution of the number of exons classified as hybrid per gene, binned by the number of internal exons in the same genes. (**B**), distribution of the number of exons classified as hybrid in genes with at least one hybrid exon, binned by the number of internal exons in the same genes. (**C**), distribution of exon position for first exons (FE) and hybrid FE-Internal (hybrid FE-Int) (left) and last exons (LE) and hybrid internal-LE (hybrid Int-LE) (right) expressed in LCL cells. Position 0 is set to the start coordinate of the most upstream first exon. Distributions were smoothed with Kernel density estimation. Percent of exons classified in 5 categories upstream and downstream of hybrid FE-Int (**D**) and hybrid internal-LE (**E**). (**F**), location of CpG islands within 300 nucleotide regions upstream of hybrid FE-Internal exons.

**Figure S8.**
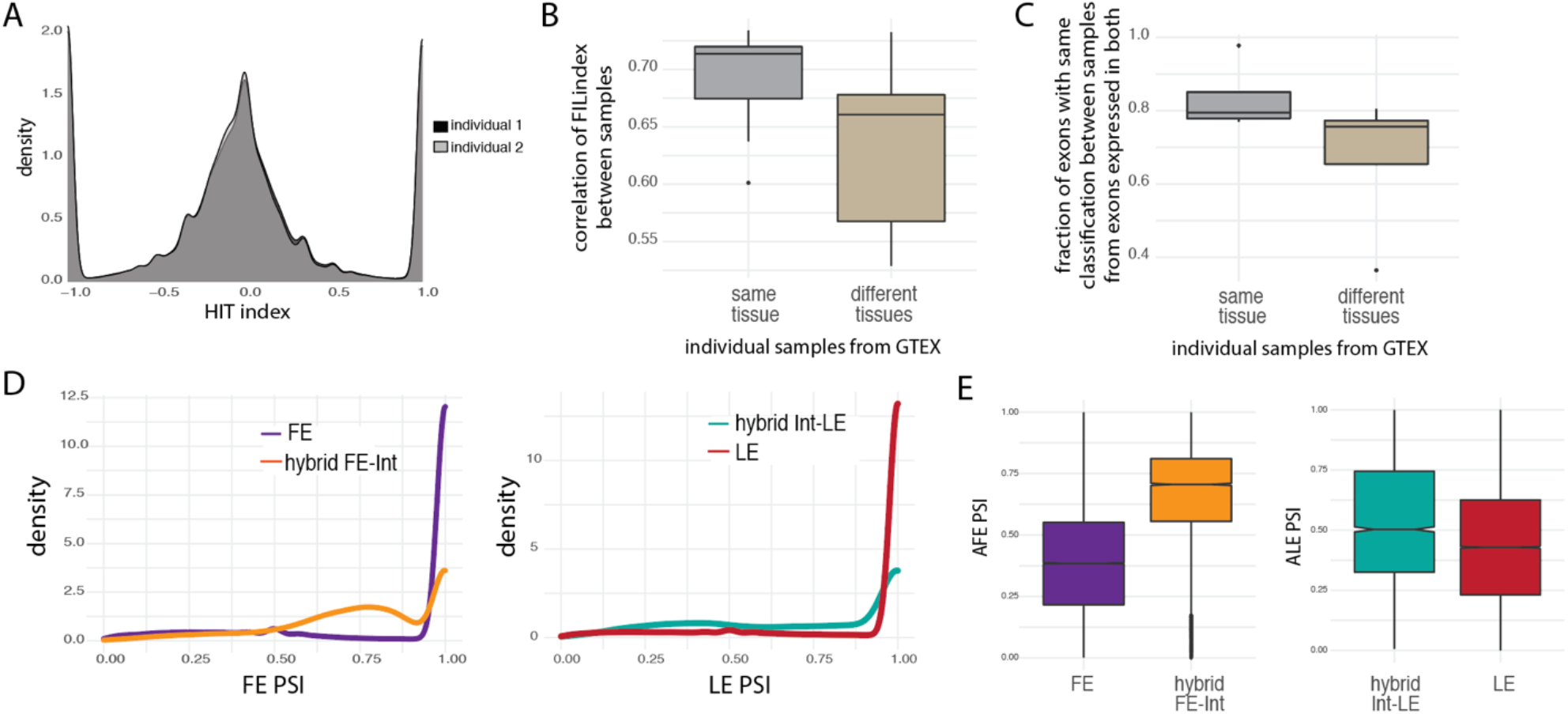
(**A**), HIT index distribution of exons expressed (> 10 junction reads) comparing data from the same tissues in two different individuals. (**B**), distribution of the spearman correlation of the HITindex between samples from different individuals considering same or different tissues. (**C**), fraction of exons classified in the same category between samples from different individuals considering same or different tissues. PSI value density (**D**) and boxplot (**E**) of alternative FE (left) and alternative LE (right) including hybrid exons across all human tissues analyzed.

**Figure S9.**
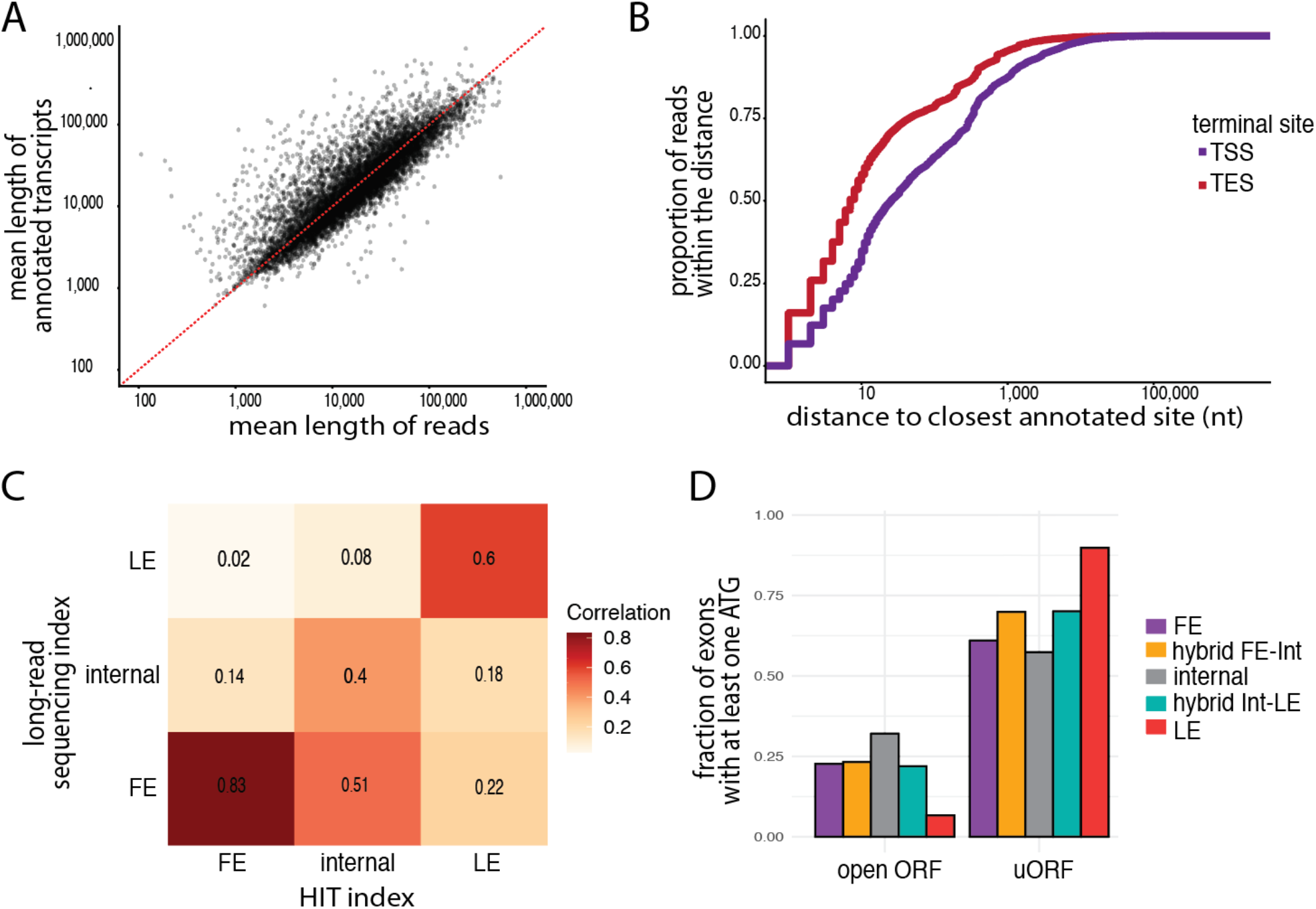
(**A**), Lengths of reads from direct mRNA sequencing (x-axis) vs. corresponding annotated transcript lengths (y-axis). (**B**), Cumulative distributions of read start and end distances from annotated TSS (*purple*) and TES (*red*) sites, respectively. (**C**), The correlation between exon classification from long-read sequencing and the HIT index pipeline.

